# A rear-engine drives adherent tissue migration *in vivo*

**DOI:** 10.1101/2021.08.03.454898

**Authors:** Naoya Yamaguchi, Ziyi Zhang, Teseo Schneider, Biran Wang, Daniele Panozzo, Holger Knaut

## Abstract

During animal embryogenesis, homeostasis and disease, tissues push and pull on their surroundings to move forward. Although the force-generating machinery is known, it is unknown how tissues exert physical stresses on their substrate to generate motion *in vivo*. Here, we identify the force transmission machinery, the substrate, and the stresses that a tissue, the zebrafish posterior lateral line primordium, generates during its migration. We find that the primordium couples actin flow through integrins to the basement membrane for forward movement. Talin/integrin-mediated coupling is required for efficient migration and its loss is partly compensated for by increased actin flow. Using Embryogram, an approach to measure stresses *in vivo*, we show that the primordium’s rear exerts high stresses, indicating that this tissue pushes itself forward with its back. This unexpected strategy likely also underlies the motion of other tissues in animals

## Main

During development, homeostasis, and disease, cells and tissues move to form organs, seal wounds and hunt pathogens^1^. To move, cells generate force and interact with their surroundings to pull and push themselves forward. Force transmission from the actomyosin network to the surroundings has been molecularly characterized and precisely measured in cultured cells^2,3^. Cells use integrin-based adhesion complexes to couple the actomyosin network inside the cells to the substrates outside the cells and pull on their surroundings with forces around 3–30 pN^4^. Since many processes are altered when cells are removed from their physiological environment and placed in culture, it is largely unclear whether cells in living animals interact with their surrounding in the same manner and pull on their substrate with similar forces. To address these questions, we used the zebrafish posterior lateral line primordium as a model. The primordium is a tissue of about 140 cells that expresses the chemokine receptor Cxcr4b. It migrates directly under the skin from behind the ear to the tip of the tail and follows a gradient of the chemokine Cxcl12a along the body of the embryo^5^.

To learn about the substrate that the primordium uses to push and pull itself forward, we inspected transverse sections of 36 hpf embryos at different locations along the primordium’s migratory route by transmission electron microscopy (TEM). Consistent with previous studies^6,7^, we found that the two-layered skin is separated from the underlying muscle by a 200 nm thick basement membrane (BM) in front of the migrating primordium (Extended Data Fig. 1a, e). At the position of the primordium, the migrating tissue separates the skin and the BM, such that the primordium’s basal side is juxtaposed to the BM while there is no BM detectable on the primordium’s apical side (Fig. 1a, Extended Data Fig. 1a–d). We confirmed these observations by inspecting the localization of the core BM component Laminin-*γ*1 tagged with superfolder GFP (LamC1-sfGFP) and expressed from the *lamC1* locus on a bacterial artificial chromosome (BAC) in live embryos (Extended Data Fig. 2a, b). During its migration the primordium wedges itself between the skin and the muscle, and pushes the LamC1-sfGFP-labeled BM towards its basal side. This separates the BM from the skin (Fig. 1b). Thus, the primordium migrates on top of a BM and directly underneath the skin. This suggests that the BM serves as the primordium’s substrate for migration.

**Fig. 1.**
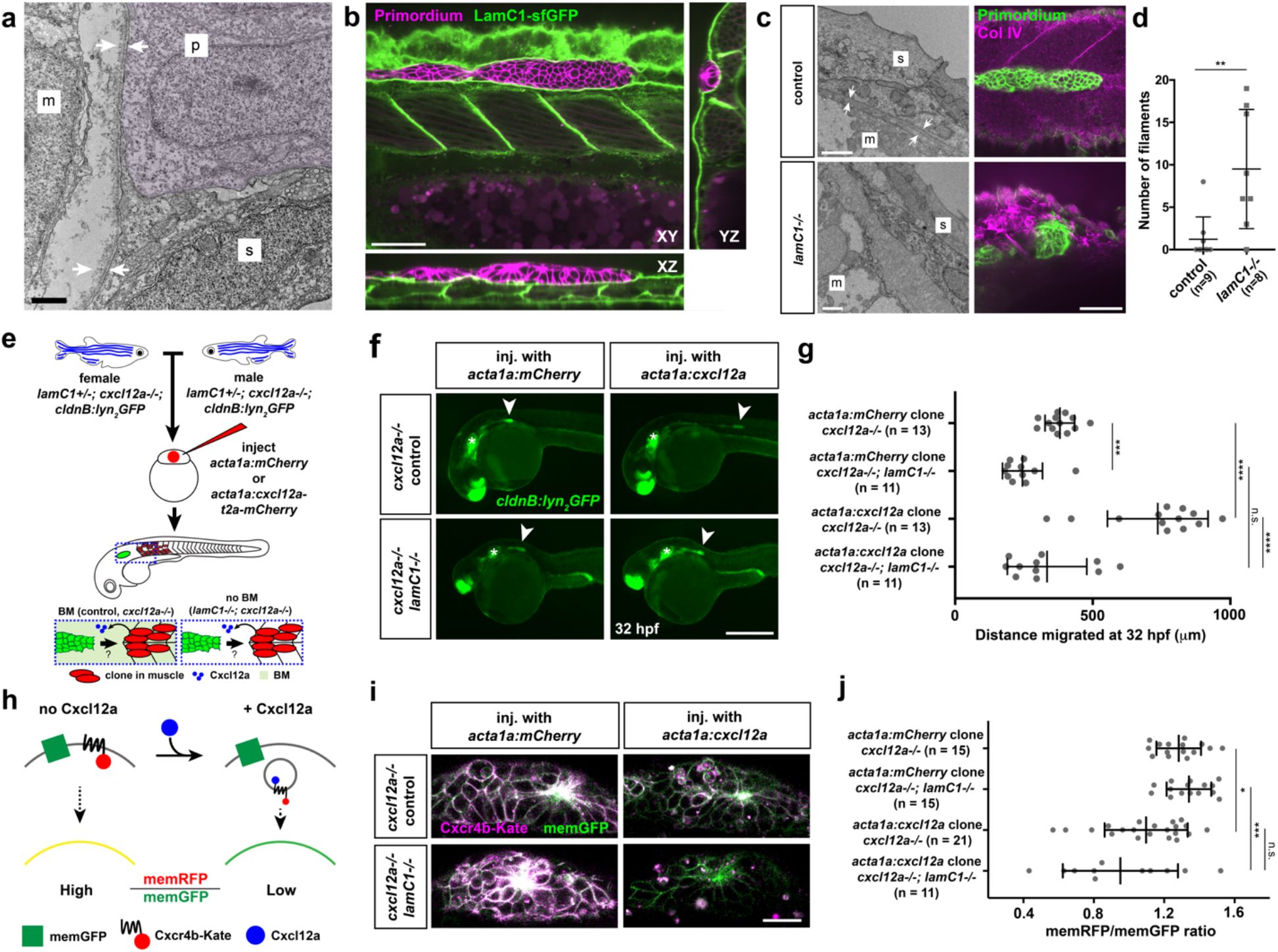
Primordium migration requires an intact basement membrane. **a**, TEM images of the skin (s), primordium (p, purple hue) and the muscle (m). White arrows indicate the BM. Scale bar = 1 *μ*m. **b**, Optical sections along the indicated planes of a z-stack of the primordium and the BM in a live 31 hpf embryo. Scale bar = 50 *μ*m. **c**, Left. TEM images of the ultrastructure of the BM between the skin (s) and the muscle (m) in control and *lamC1*-/-embryos. White arrows indicate the BM. Scale bar = 2 *μ*m. Right. Antibody staining against Collagen IV in control and *lamC1*-/- embryos. Scale bar = 50 *μ*m. **d**, Quantification of Collagen IV filaments in control and *lamC1*-/- embryos. **: p<0.001 (Mann-Whitney test). **e**, Strategy to express Cxcl12a in a few muscle cells in *lamC1*-/- embryos and siblings. **f**, Images of the migrating primordium in *cxcl12a*-/- and *cxcl12a*-/-; *lamC1*-/- 32 hpf embryos with clones in the muscle of the trunk that express mCherry (not shown) or Cxcl12a together with mCherry (not shown). Asterisks indicate the ear and arrowheads the primordium. Scale bar = 0.5 mm. **g**, Quantification of the distance migrated by the primordium in the indicated experimental conditions at 32 hpf. ****: p<0.0001, n.s: p>0.05 (Mann-Whitney test). **h**, Principle of the Cxcl12a sensor. **i**, Images of the Cxcl12a sensor in primordia of *cxcl12a*-/- and *cxcl12a*-/-; *lamC1*-/- live embryos with clones in the muscle of the trunk that express mCherry or Cxcl12a. Scale bar = 20 *μ*m. **j**, Quantification of the Cxcr4b-Kate-to-memGFP ratio in the primordia of embryos described in (**i**). *: p<0.05, ***: p<0.001, n.s: p>0.05 (one way ANOVA followed by Holm-Sidak’s multiple comparison test). Note, controls are *lamc1+/+* and *lamc1−/+* embryos.

If the primordium uses the BM as its substrate to push itself forward, the primordium should exert stresses (force per area) on the BM and deform the BM. Such traction stresses have been measured for migrating cells in culture by imaging the displacement of fluorescent beads embedded in elastic surfaces or matrices and the bending of flexible cantilevers—collectively referred to as traction force microscopy^8–16^. To extend traction force microscopy to living embryos, we created optical landmarks on the BM and assessed how these landmarks are displaced as the primordium moves across them (Fig. 2a). Using a laser, we locally bleached LamC1-sfGFP in an approximately cylindrical volume in the BM (Extended Data Fig. 2h, i, Video 1). Little LamC1-sfGFP diffused back into the bleached cylinder (*t*_1/2_=12.3 min, mobile fraction = 20.1%, Extended Data Fig. 2j, k), and bleached cylinders remained clearly demarcated for two hours after photo-bleaching (Video 2) while untagged, extracellular mCherry filled bleached cylinders rapidly after bleaching (*t*_1/2_=1.1 sec, mobile fraction = 62.6%, Extended Data Fig. 2j). This indicated that bleaching LamC1-sfGFP is a suitable approach to place local marks on the BM and monitor the BM’s deformation over time. We therefore bleached a hexagonal pattern of marks onto the LamC1-sfGFP-labeled BM with a distance of 6 *μ*m between individual marks and a mark diameter of 2 *μ*m (Extended Data Fig. 2i). We generated this pattern directly in front of the primordium and across its migratory route, and recorded the position of the marks as the primordium migrated across this pattern (Video 2). To reconstruct the stresses from the displacement of the marks on the BM by the migrating primordium, we developed the analysis pipeline Embryogram (Fig. 2e) which was inspired by the Cellogram algorithm^17^ developed for tracking quantum dots in confocal monocrystalline arrays. Embryogram identifies the bleached cylinders in the first frame of the time lapse by fitting cylindrical volumes using numerical methods that combine coarse sampling of the solution space with local fitting using the quasi Newton algorithm. This assigns a point in space to each mark in the image frame. To connect the points to obtain a triangular mesh, we start from a regular triangular mesh and fit it over the marker positions using a variant of the iterative closest point algorithm. This fitting is only done for the first frame of the time lapse, and the resulting triangular mesh is an approximation of the pattern of the marks on the BM. This mesh is then deformed to follow the marks on the BM in the subsequent frames of the time lapse. For each subsequent frame, we use 3-dimensional optical flow to compute a smooth and coarse map between the current and the previous frame, which is applied to the mesh. The location of each mark is then fine-tuned over the new image frame using the same algorithm used to fit the marks in the first frame of the time lapse. This procedure is repeated until the end of the time lapse, leading to a time-sequence of triangular meshes that captures the deformation of the BM. To correct for rigid motions of the sample during imaging, we compute the best-fitting rototranslation between the mesh at every frame and the initial mesh over a subset of marks that are not deformed by the migrating primordium (Extended Data Fig. 3a). The time-varying mesh is used to compute displacements between the marks. To convert the displacements into stresses, we fill an axis-aligned box that contains the sample with a volumetric tetrahedral mesh. This mesh contains (and conforms to) the surface mesh approximating the BM in the first frame of the time lapse. The stresses are then computed solving an elastic deformation of the volumetric mesh, while keeping the boundary of the mesh fixed, and applying the displacements extracted by our tracking algorithm as Dirichlet boundary conditions for the nodes corresponding to the BM (for details see Supplemental Note). For this conversion, we determined the stiffness, or Young’s modulus, of the BM. The Young’s modulus describes the relationship between tensile stress and axial strain. We removed the skin above the BM and measured the BM stiffness by atomic force microscopy (AFM) (Fig. 2a–c, Extended Data Fig. 3b–h). In agreement with previous *in vivo* studies assessing stiffness at the micron-scale^18,19^, this analysis yielded a Young’s modulus for the BM of 566 ± 355 Pa (mean and SD) which was reduced to 321 ± 158 Pa (mean and SD) after collagenase treatment (Fig. 2d, Extended Data Fig. 3b, e), a value probably reflecting the stiffness of the muscle that underlies the 200 nm thick BM (Extended Data Fig. 3c, g). Importantly, spontaneous twitches of skin cells contract the underlying BM. This causes the BM to buckle and wrinkle akin to the distortions of the substrate observed around cultured cells^20^ and in animals^21^. The wrinkles form and disappear in less than 2 min and tracking the optical marks indicates that the wrinkles do not cause lasting deformation of the BM (Extended Data Fig. 4a–e, Video 3). Similarly, repeated probing of the BM at the same location in deskinned embryos by AFM did not alter the stiffness measurements (Extended Data Fig. 3f). These observations suggest that the BM undergoes non-plastic deformations in response to endogenous and external stresses and can be approximated by a linear stress-strain relationship.

**Fig. 2.**
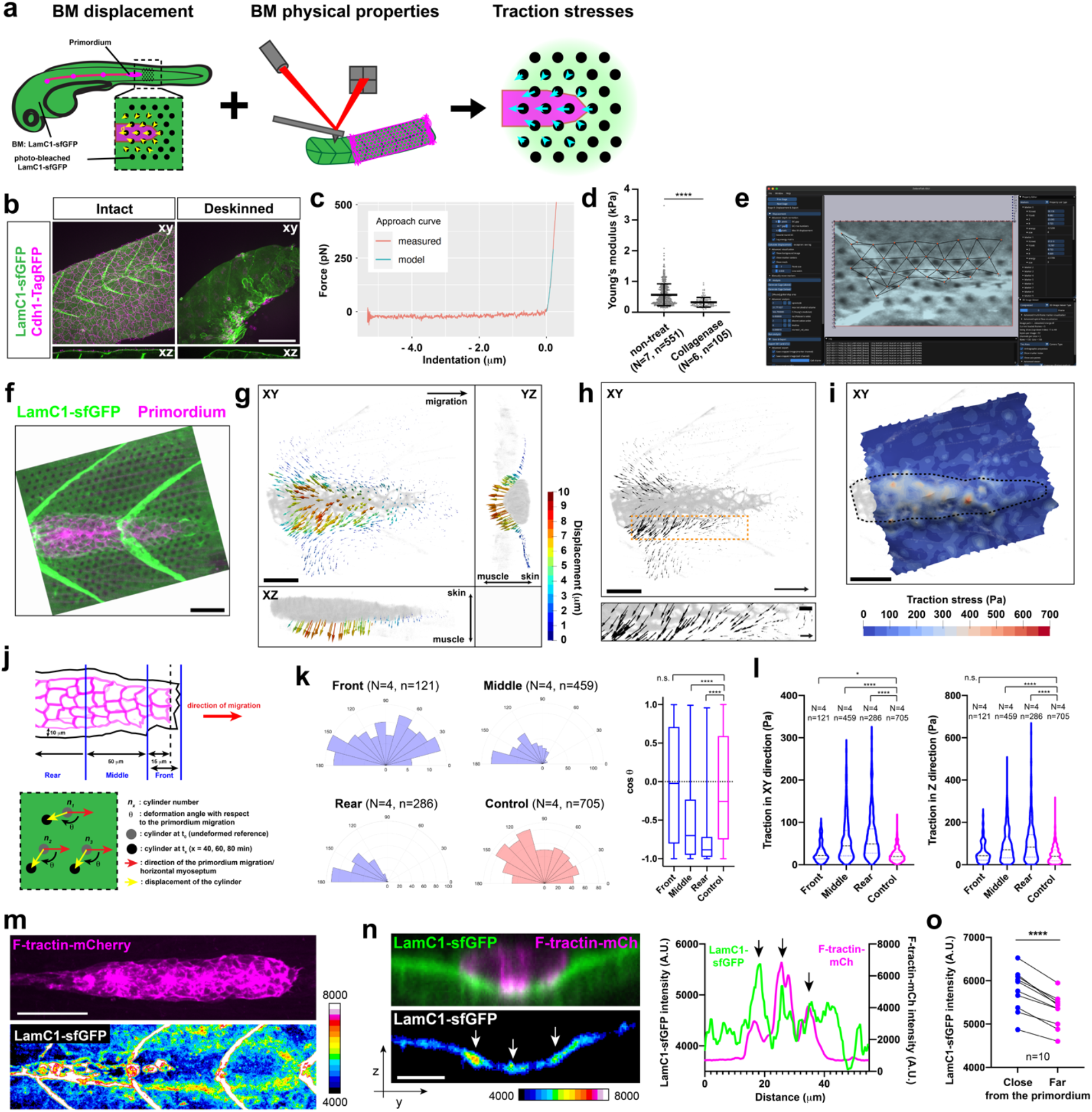
Stress measurements indicate that the primordium exerts high stresses in its rear. **a**, Strategy to measure stresses *in vivo*. **b**, 28 hpf embryos with labeled skin and BM before and after surgical skin removal. Images are maximum-projected z-stacks. Scale bar = 100 *μ*m. **c**, Representative force curve of BM stiffness measurement by AFM. Fit of the first 200 nm after the contact point at 0 *μ*m to the Hertz model is indicated in cyan. Only the approach part of the force curve is shown. **d**, Quantification of the BM stiffness with and without Collagenase treatment. ****: p<0.0001 (Mann-Whitney test). **e**, Image of Embryogram application user interface. **f**, Maximum-projected z-stack at 90 min taken from Video 2. Scale bar = 25 *μ*m. **g**, BM displacement by the primordium shown as a vector field along the X-, Y- and Z-axes. The primordium is indicated in gray. The displacement vector magnitude is indicated as a color map. Note that the XY-view is shown from the basal side of the primordium. Image taken from Video 4 at 90 min. Gray scale bar = 25 *μ*m. **h**, BM displacement in the XY-plane by the primordium shown as a vector field. The primordium is indicated in gray. Note that the displacement field in the XY-plane is shown from the basal side of the primordium. Image taken from Video 4 at 90 min. Scale bar = 25 *μ*m. Bottom panel is a magnification of the region outlined with an orange rectangle. Scale bars = 25 *μ*m (top) and 5 *μ*m (bottom), arrow = 50 *μ*m (top) and 10 *μ*m (bottom). **i**, Distribution of traction stress magnitudes on the BM. The stress magnitude is indicated as a color map. Note that the stress distribution is shown from the basal side of the primordium. Image taken from Video 4 at 90 min. Scale bar = 25 *μ*m. **j**, Schematic of the approach to quantify the traction stresses and BM displacement vector direction along the front-rear axis of the primordium. **k**, BM displacement vector angles with respect to the migration direction (0°). ****: p<0.0001 and n.s.: p>0.05 (Mann-Whitney test). N = number of embryos, n = number of bleached cylinders analyzed. **l**, Pooled traction stresses exerted by the primordium on the BM. Magnitude in XY-plane (left) and along the Z-axis (right) are shown. ****: p<0.0001 and n.s.: p>0.05 (Mann-Whitney test). N = number of embryos, n = number of bleached cylinders analyzed. **m**, F-tractin-mCherry distribution in the primordium (top) and LamC1-sfGFP around the primordium (bottom) in a 32 hpf embryo. Note that LamC1-GFP fluorescence intensity is pseudo-colored as a heat map. The image is a maximum-projected z-stack. Heat map ranges from 4000 to 8000. Scale bar = 50 *μ*m. **n**, Transverse section through the F-tractin-mCherry-expressing primordium and the underlying LamC1-sfGFP-labeled BM (left top). Corresponding image showing the LamC1-sfGFP fluorescence intensity as a heat map (left bottom). Arrows indicate apposed clusters of F-tractin-mCherry and LamC1-sfGFP. Images are single sections along the YZ-plane of a z-stack. Scale bar = 10 *μ*m. Fluorescent intensity profiles of F-tractin-mCherry and LamC1-sfGFP of image shown in left along the Y-axis (right). Arrows indicate the position of the apposed clusters of F-tractin-mCherry and LamC1-sfGFP indicated by arrows in left. **o**, Quantification of the LamC1-sfGFP intensity within 3 *μ*m-wide bands around the perimeter of the primordium (left) and at a distance of 6 *μ*m from the primordium’s perimeter (right). ****: p< 0.0001 (Paired t-test).

Using the Embryogram pipeline together with the stiffness measurements, we find that the migrating primordium pulls the BM slightly sideways and backwards at its tip while also pushing it downward. Towards its rear, this pattern is more pronounced and along its sides the primordium pushes more strongly backwards than in its front (Fig. 2f–h, j, k, Video 4). The traction stresses (the forces against the plane of the BM) reflect this displacement pattern. In the front, the mean traction stresses average 28 ± 22 Pa, increase to 58 ± 49 Pa and 64 ± 51 Pa (mean and SD) in the middle and rear of the primordium, respectively, with higher traction stresses exerted preferentially along the sides of the primordium peaking at 600 Pa (Fig. 2i, j, l, Video 4). Similarly, the primordium generates high, mostly rearward-pointing stresses (stresses extracted in the direction of migration) along the sides and towards its rear (Extended Data Fig. 4f, where it also exerts the highest shear stresses on the BM (Extended Data Fig. 4g). Although the primordium moves at fairly constant speed^22^, it does not exert constant stresses on its substrate (Video 4), suggesting that its forward motion is the result of the average of the fluctuating stresses across the tissue. At the position where the primordium has passed, the BM returned to its original shape, indicating that the BM is not irreversibly deformed by the primordium (Video 4). Also, we did not observe such BM displacements and stresses in controls in which we blocked primordium migration by ubiquitous over-expression of the primordium’s attractive guidance cue Cxcl12a (Fig. 2k, l, Extended Data Fig. 4h–m, Video 4). The observed stress distribution was also reflected in the wrinkling of the BM along the sides of the primordium with LamC1-sfGFP forming local clusters (Fig. 2m–o). Together, this indicates that the primordium moves in a continuous breaststroke-like manner with its front and middle pushing the BM sideways and downward while the cells on its side and rear push sideways and strongly backwards, consistent with theoretical predictions for adherent cell migration^23^.

Since the BM serves as a substrate for the primordium the BM should also be required for the migration of the primordium. To test this prediction, we analyzed the migration of the primordium in *lamC1* mutant embryos. In such embryos, the BM is disrupted or missing^7^, and inspection of the Collagen IV network and the BM confirms this observation (Fig. 1c, d). Since the lack of LamC1 also impairs the formation of the Cxcl12a-secreting stripe of cells that guides the primordium (Extended Data Fig. 2e), we assessed the ability of the primordium to migrate in *lamC1* mutant embryos by generating an ectopic Cxcl12a source in the trunk muscles. The initial location of the primordium is not affected in *lamC1* mutant embryos (Extended Data Fig. 2f, g). For this analysis, we also removed endogenous Cxcl12a to avoid competition between endogenous and ectopic chemokine sources (Fig. 1e). While local secretion of Cxcl12a from the muscle of *cxcl12a* mutant control embryos attracted the primordium, Cxcl12a failed to restore directed migration towards the chemokine sources in *lamC1; cxcl12a* double mutant embryos (Fig. 1f, g). Ectopic Cxcl12a triggered the internalization of its receptor Cxcr4b in primordia of *lamC1; cxcl12a* double mutant embryos and *cxcl12a* mutant control embryos, indicating that the diffusion and presentation of Cxcl12a is not impaired in the absence of LamC1 and the BM (Fig. 1h–j). Thus, an intact BM is required for directed primordium migration, confirming that the primordium uses the BM as its substrate for migration.

To push, pull, and exert stresses on the BM, the primordium needs to adhere to the BM. Molecularly, cells can adhere to the BM through focal adhesions. Two core components of these large protein complexes are integrins and talins^24^. Integrins bind to specific BM components on the outside of the cell and—through talins and other adaptors—to the actin network inside the cell. To test whether the primordium uses focal adhesions to interact with the BM, we first identified the *β*-integrins and talins that the primordium expresses. Of the twelve *β*-integrins and three talins in zebrafish, *integrin-β1b* (*itgb1b*) and *talin1* (*tln1*) were expressed throughout the primordium (Extended Data Fig. 5a, 6a). We therefore tagged *itgb1b* and *tln1* with superfolder GFP and YPet at the endogenous locus and on a BAC transgene, respectively (Extended Data Fig. 5h, i, 6c, d). Itgb1b-sfGFP and Tln1-YPet recapitulated the endogenous expression pattern and restored viability when placed in the respective mutant background (*itgb1b-sfGFP/-* and *tln:tln1-YPet; tln1-/-*) (Fig. 4d, Extended Data Fig. 5i, 6d, data not shown). While Itgb1b-sfGFP and Tln1-YPet were enriched at the myotendinous junctions of the muscle, Itgb1b-sfGFP localized fairly uniformly on the membranes of the primordium cells (Fig. 3a, Video 5), and Tln1-YPet was mostly cytoplasmic in the cells of the primordium (Fig. 3b, Video 5). Since Itgb1b-sfGFP and Tln1-YPet are also expressed by the surrounding skin and muscle, this expression could mask protein clustering on the membranes of the primordium cells. To circumvent this potential problem, we generated embryos in which only a few cells in the primordium expressed Itgb1b-sfGFP or Tln1-YPet together with membrane-tethered mCherry by blastomere transplantation (Fig. 3c). This analysis revealed that Itgb1b-sfGFP and Tln1-YPet formed short-lived clusters with a lifetime of less than 2 minutes on the basal sides of the cells in the primordium, often within the basal protrusions (Extended Data Fig. 7a, b). Since talin links integrin to F-actin^24^, we asked whether clustered Itgb1b and Tln1 co-localized with F-actin. Chimeric analysis showed that this is the case; Itgb1b-sfGFP and Tln1-YPet clusters co-localized with F-tractin-labeled F-actin on the basal sides of primordium cells (Fig. 3d–f, Video 6). Control experiments showed that Itgb1b-sfGFP and, to a lesser degree, Tln1-YPet also co-localized with membrane-tethered mCherry, as expected for a transmembrane protein and a cytosolic protein, respectively (Extended Data Fig. 7c, d). Thus, the primordium cells form small, transient integrin/talin/F-actin clusters on their basal sides. We corroborated the transient nature of the integrin clusters by measuring the mobility of integrin in the membrane through FRAP. Ligated integrin couples to the actin network and diffuses more slowly in the membrane than unligated integrin^25,26^. The mobility of integrin is therefore a measure of the degree of ligated integrins interacting with actin. Consistent with integrin function in muscle, the mobility of Itgb1b-sfGFP at the myotendinous junction is low (*t*_1/2_=17.6 sec, mobile fraction = 20.1%) and increases when blocking ROCK-mediated actin network contractions (*t*_1/2_=14.9 sec, mobile fraction = 27.1%) (Extended Data Fig. 8f–h). In contrast, the mobility of Itgb1b-sfGFP is high in the cells of the primordium (*t*_1/2_=11.4 sec, mobile fraction = 33.6%, Extended Data Fig. 8i–k), supporting the idea that integrin interacts with the actin network only transiently in this migrating tissue. In contrast to migrating cells *in vitro*^27,28^, these observations suggest that the primordium cells do not form long-lived focal adhesions but rather transient integrin clusters. When placed on a Laminin-coated surface, primordium cells formed large integrin and talin clusters along F-actin cables as observed in cultured cells^27,28^ (Extended Data Fig. 7e–g), indicating that primordium cells can form focal adhesions and stress fibers *ex vivo* but do not do so *in vivo*.

**Fig. 3.**
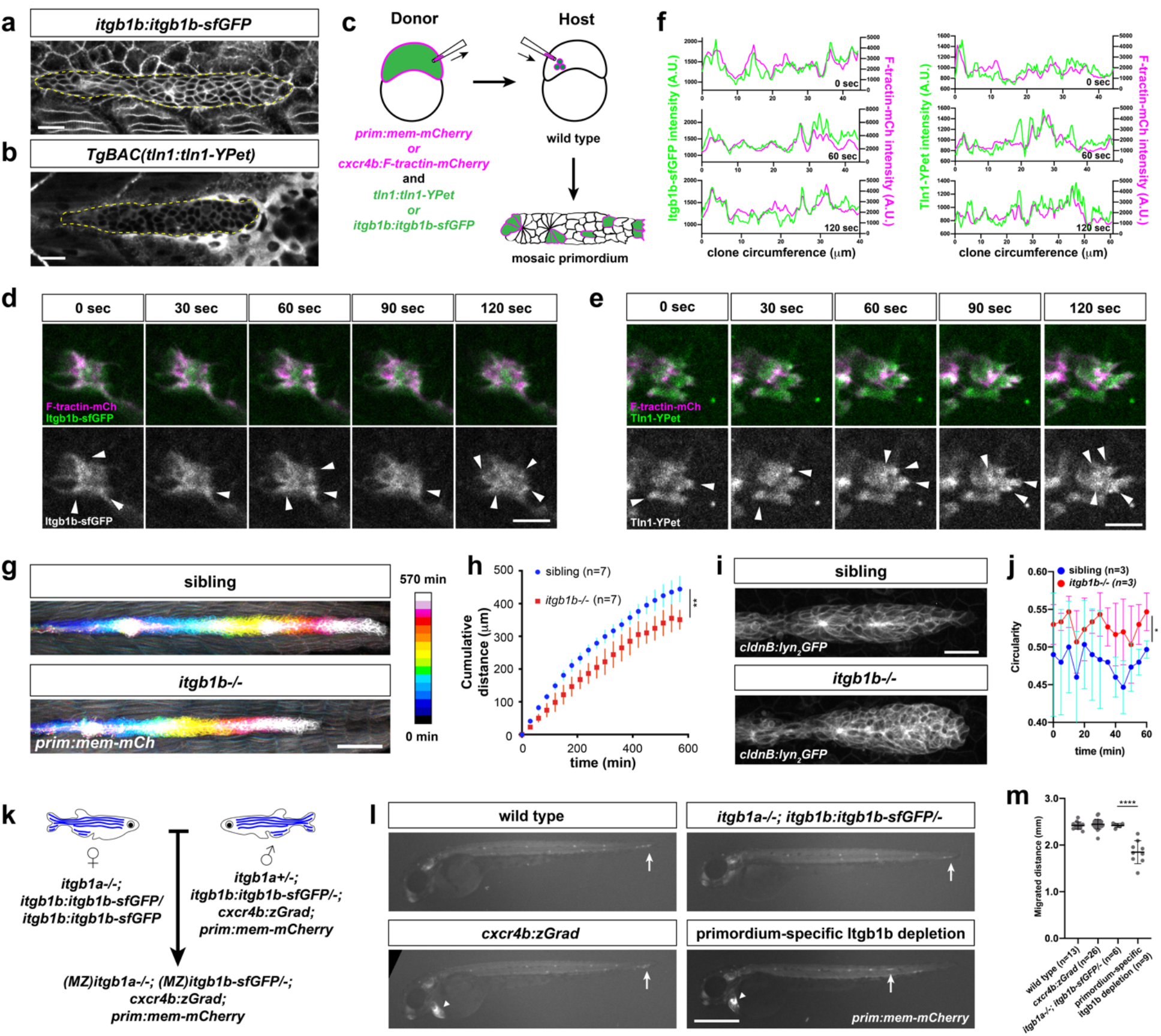
*β*1-Integrin forms clusters at the basal sides of the cells in the primordium and is required for efficient migration. **a**, Expression of Itgb1b-sfGFP from the endogenous locus in a 33 hpf embryo. The image is a single slice from a z-stack. The primordium is outlined by a dotted, yellow line. Scale bar = 20 *μ*m. **b**, Expression of Tln1-YPet from the *tln1:tln1-YPet* BAC transgene in a 33 hpf embryo. The image is a single slice from a z-stack. The primordium is outlined by a dotted, yellow line. Scale bar = 20 *μ*m. **c**, Schematic of blastomere transplantation experiments. **d**, Images of Itgb1b-sfGFP and F-tractin-mCherry localization at the basal side of a clone in the primordium over time. Images are single slices from a time-lapse movie (Video 6). Arrowheads indicate Itgb1b-sfGFP/F-tractin-mCherry clusters. Scale bar = 10 *μ*m. **e**, Images of Tln1-YPet and F-tractin-mCherry localization at the basal side of a clone in the primordium over time. Images are single slices from a time-lapse movie (Video 6). Arrowheads indicate Tln1-YPet/F-tractin-mCherry clusters. Scale bar = 10 *μ*m. **f**, Intensity profiles of Itgb1b-sfGFP (left) and Tln1-YPet (right) with F-tractin-mCherry along the perimeter on the basal sides of the clones shown in (**d**) and (**e**) over time. **g**, Primordium migration in wild-type and *itgb1b-/-* embryos. Images are maximum-projected z-stacks that were temporally color coded from 0 to 570 min. Scale bar = 100 *μ*m. **h**, Quantification of the cumulative primordium migration distance in control (blue) and *itgb1b-/-* embryos (red). Mean (dots) and SD (bars) are indicated. **: p<0.01 (t-test at the end point). **i**, Images of primordia in wild-type and the *itgb1b-/-* embryos. Images are maximum-projected z-stacks. Scale bar = 25 *μ*m. **j**, Quantification of primordium circularity in control (blue) and *itgb1b-/-* embryos (red). Mean (dots) and SD (bars) are indicated. *: p<0.05 (t-test at the end point). **k**, Crossing scheme to generate embryos with primordium-specific depletion of Itgb1b-sfGFP. **l**, Degree of primordium migration in 48 hpf embryos of indicated genotypes. Primordium-specific Itgb1 depletion refers to genotype shown in (**k**). Arrows indicate primordia, arrowheads indicate the transgene marker for *cxcr4b:zGrad*. Scale bar = 500 *μ*m. **m**, Quantification of the primordium migration distance of embryos at 48 hpf shown in (**l**). ****: p<0.0001 (one way ANOVA followed by Tukey’s multiple comparison test).

**Fig. 4.**
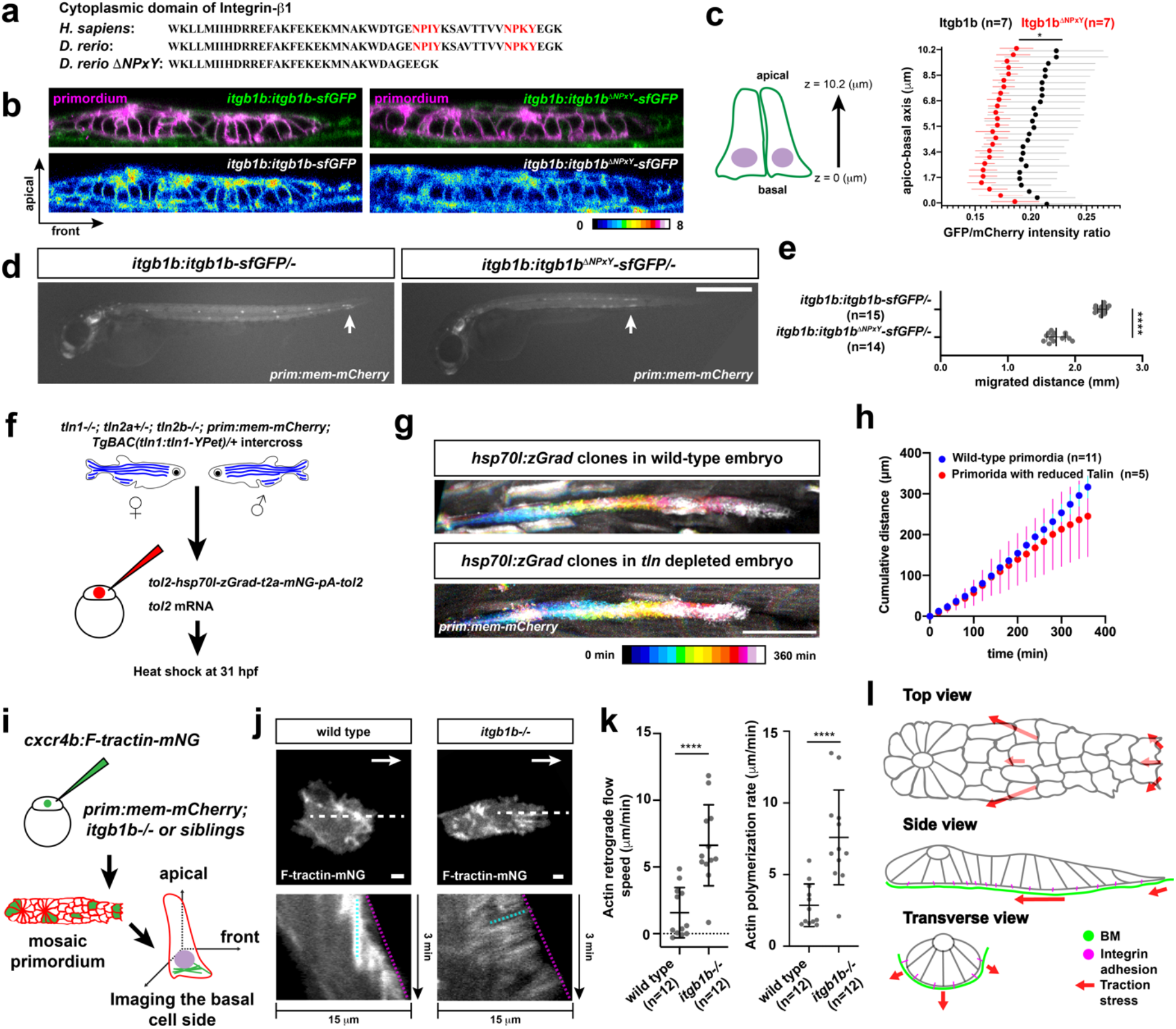
*β*1-Integrin couples cell-substrate adhesion to actin flow in the primordium. **a**, Alignment of amino acid sequences of the human and zebrafish Integrin-*β*1’s cytoplasmic domains, and the zebrafish Itgb1b^*ΔNPxY*^ mutant allele. The two NPxY motives are indicated in red. **b**, Localization of Itgb1b-sfGFP (left) and Itgb1b^*ΔNPxY*^-sfGFP (right) in the primordium. Images are single transverse section from z-stacks. Scale bar = 50 *μ*m. **c**, Quantification of the Itgb1b-sfGFP and Itgb1b^*ΔNPxY*^-sfGFP distribution on the membranes of primordium cells along the apicobasal axis normalized to the fluorescence intensity of membrane-tethered mCherry. *: p<0.05 (Welch’s t-test). **d**, Images of the distance migrated by the primordium in *itgb1b:itgb1b-sfGFP/-* (left) and the *itgb1b:itgb1b*^*ΔNPxY*^*-sfGFP/-* embryos (right) at 48 hpf. Arrows indicate primordia. Scale bar = 0.5 mm. **e**, Quantification of the primordium migration distance at 48 hpf. ****: p<0.0001 (Mann-Whitney test). **f**, Experimental strategy to generate cell clones in the primordium with heat shock inducible zGrad expression. **g**, Migration of primordia with cell clones expressing zGrad from a heat shock inducible promoter in control embryos (top) and embryos with reduced Talin activity (bottom). Images are sum-projected z-stacks that were temporally color coded from 0 to 360 min. Scale bar = 100 *μ*m. **h**, Quantification of the cumulative primordium migration distance in control embryos (blue) and embryos with *hsp70l:zGrad* cell clones in the primordium (red). Mean (dots) and SD (bars) are indicated. **i**, Experimental design to assess actin flow. **j**, Images of F-tractin-mNeonGreen localization at the basal sides of wild-type and *itgb1b-/-* primordium cells (top). White arrows indicate the direction of migration. Scale bar = 2 *μ*m. Images are single optical sections from Video 10. Kymographs of Video 10 along the dotted line indicated in top images (bottom). The dotted cyan and magenta lines indicate the rates of actin flow and protrusion, respectively. **k**, Quantification of the rates of actin flow (left) and actin polymerization (right). ****: p<0.0001 (Mann-Whitney test). **l**, Model of Primordium motility. The cells in the primordium transduce force across integrins to the BM and exert small, and mostly sideways directed stresses in the front, and larger, mostly rearward directed stresses in the back. Combined these stresses push the primordium forward.

Next, we asked whether Integrin and Talin function are required for primordium migration. Since *itgb1b* and *tln1*—and possibly *itgb1a*, *tln2a* and *tln2b*—are expressed in the primordium, we generated mutants in these five genes (Extended Data Fig. 5b, 6b). Phenotypic analysis showed that the primordium was less elongated and migrated more slowly in *itgb1b* mutant embryos than in wild-type controls (Fig. 3g–j, Extended Data Fig. 5c–g, Video 7). Since *itgb1a*−/+; *itgb1b*-/- and *itgb1a*-/-; *itgb1b*-/- embryos have severe morphogenesis defects (Extended Data Fig. 5c), we could not assess primordium migration in these genetic scenarios. Instead we assessed whether Itgb1b is required within the tissue for migration by depleting Itgb1b-sfGFP in the primordium of *itgb1a*-/-; *itgb1b-sfGFP*/-embryos using zGrad, a degron system based on an anti-GFP nanobody^29^ (Extended Data Fig. 9a–d), driven from the *cxcr4b* promoter that expresses in the migrating primordium (Fig. 3k). In such embryos, primordium migration was impaired to almost the same degree as in *itgb1b* mutant embryos (Fig. 3l, m), indicating that Itgb1b is required within the primordium for migration with minor contribution from Itgb1a.

In *talin* single, double and triple mutant embryos, the primordium migrated normally and morphogenesis was mostly unaffected—the heart was smaller and oedemic, and the body length was slightly reduced compared to wild-type embryos (Extended Data Fig. 6e, f). A possible explanation for the surprisingly mild defects in *tln1*-/-; *tln2a*-/-; *tln2b*-/-embryos could be the maternal contribution of *talin* mRNA and protein to the embryo. To address this possibility, we generated zygotic (Z) *Z tln1*-/-; *Z tln2a*-/-; *Z tln2b*-/- embryos which also lacked the maternal contribution (M) of *M tln1* and *M tln2b*. Such embryos were obtained by in-crossing *tln1*-/-; *tln2a*−/+; *tln2b*-/-; *tln1:tln1-YPet*; *prim:mem-mCherry* fish, and injecting the embryos with *zGrad* mRNA (Extended Data Fig. 6g). *tln1:tln1-YPet* rescues the lethality of *tln1* mutants, *prim:mem-mCherry* marks the primordium, and zGrad protein expressed from *zGrad* mRNA degrades maternally deposited Tln1-YPet protein. Because *MZ tln1*-/-; *Z tln2a*-/-; *MZ tln2b*-/- embryos had somitogenesis defects and disrupted *cxcl12a* expression along the migratory route of the primordium (Extended Data Fig. 6h–j), we could not assess the role of Talin in primordium migration. We therefore depleted Tln1-YPet in a few primordium cells after the beginning of migration by injecting *hsp70l:zGrad-t2a-mNeonGreen* DNA in one-cell stage *MZ tln1*-/-; *Z tln2a*-/- or *Z tln2a*+/−; *MZ tln2b*-/-; *tln1:tln1-YPet*; *prim:mem-mCherry* embryos and heat-shocking the embryos at 31 hpf (Fig. 4f). The injected *hsp70l:zGrad-t2a-mNeonGreen* DNA is inherited only by a few cells in the primordium which—upon heat shock—will express zGrad together with mNeonGreen (mNeonGreen is not degraded by zGrad, Extended Data Fig. 9b), and deplete Tln1-YPet (Extended Data Fig. 9e, f). This analysis showed that depleting most Talin activity in a few primordium cells slowed the migration of the primordium compared to the migration of primordia in control embryos (Fig. 4g, h, Video 8). We confirmed this observation by placing cells with strongly reduced Talin activity in the front part of migrating primordia by blastomere transplantation (Extended Data Fig. 6k). Compared to controls, such primordia also migrated slower (Extended Data Fig. 6l–n, Video 9). Consistent with its role in bridging integrin to the actin network, this indicates that talin is required for proper migration of the primordium.

We corroborated this idea further by disrupting the interaction between integrin and talin. Talin, and other cytosolic partners of integrin, bind to the NPxY motives in integrin’s cytoplasmic tail^24^. We therefore deleted the two NPxY motives in Itgb1b and tagged Itgb1b^*ΔNPxY*^ with superfolder GFP at its endogenous locus (Fig. 4a, Extended Data Fig. 5j). Compared to Itgb1b-sfGFP, Itgb1b^*ΔNPxY*^-sfGFP localized less to the apical and basal sides of the cells in the primordium and its levels were reduced by 16% (Fig.4b, c). This was also observed for the myotendinous junctions (Extended Data Fig. 8a, b). Consistent with its more uniform distribution, Itgb1b^*ΔNPxY*^-sfGFP was more mobile (*t*_1/2_=17.2 sec, mobile fraction = 32.8%) than Itgb1b-sfGFP in the membrane (*t*_1/2_=18.7 sec, mobile fraction = 17.4%) (Extended Data Fig. 8c–e). Similar to the global and tissue-specific loss of Itgb1b, Itgb1b^*ΔNPxY*^-sfGFP also failed to support efficient primordium migration in *itgb1b*^*ΔNPxY*^*-sfGFP*/-embryos compared to *itgb1b-sfGFP*/-control embryos (Fig. 4d, e). This indicates that the integrin/talin complex is important for the primordium to move along its migratory route.

In migrating cells, the integrin-talin complex can couple F-actin flow inside the cell to the BM outside the cell and transduce force. When engaged, the integrin-talin complex slows F-actin flow towards the rear of the cell. When disengaged, the retrograde flow of F-actin is faster^30^. To test whether the cells in the primordium use this clutch-like mechanism, we measured the speed of F-actin flow on the basal sides of cells in the primordium of wild-type and *itgb1b* mutant embryos by labeling F-actin with F-tractin-mNeonGreen in a few cells of the primordium (Fig. 4i). This analysis showed that F-actin was concentrated in the front of wild-type cells and its flow was halted or slow towards the cells’ rear with a mean speed of 1.5 *μ*m/min (Fig. 4j, k, Video 10). Conversely, F-actin formed radial cables in *itgb1b* mutant cells—similar to talin-depleted cells in culture^31^—and its mean speed of flow increased to 6.6 *μ*m/min (Fig. 4j, k, Video 10). The actin polymerization rate—the sum of the actin flow rate and the rate of membrane protrusion—also increased from 2.8 *μ*m/min in wild-type control cells to 7.6 *μ*m/min in *itgb1b* mutant cells (Fig. 4k, Extended Data Fig. 10a, b). Thus, integrins couple force for primordium motility and, similar to the situation in cultured dendritic cells and macrophages^32^, increase actin polymerization in response to decreased force coupling to the substrate to maintain forward movement—likely by coupling through other integrins and unspecific adhesion—but in contrast to the situation in dendritic cells fail to sustain wild-type speeds.

Together, this work elucidates how a tissue moves through a live animal. It provides three major insights. First, primordium cells link the force-generating actomyosin network to the BM through integrin clusters on their basal sides. These integrin clusters are less than 2 *μ*m in size and form and disassemble in less than 2 minutes. This is in contrast to the larger and longer-lived focal adhesions that migrating cells in culture use to pull themselves forward^27,28^, and more reminiscent of nascent adhesions that form at the edge of protrusions and underneath spreading cells in culture^33–36^, suggesting that tissues in animals rely on transient rather than prolonged cell-substrate interactions for movement, possibly to accommodate and adapt to a complex and rapidly changing environment. Second, the primordium cells pull and deform the basement membrane on their outside with a maximal stress close to 1 kPa. This is comparable to the average stress that migrating cells in culture exert on their substrates^8–16^, suggesting that stresses of around 1 kPa are inherent to adherent migration in simplified and physiological scenarios, possibly because these stress magnitudes reflect optimal force transmission from the actomyosin network to the outside. Consistent with this notion, retrograde actin flow is slowed or stalled in primordium cells, suggesting that most of the flow is converted into forward movement. Third, the primordium moves similar to a continuous breaststroke by pushing the BM downward, sideways and backwards. Counterintuitively, it generates most of the pulling along its sides in the rear (Fig.4l). Although surprising, this rear-engine design might, in part, be a consequence of collective force generation by the higher number of cells in the primordium’s rear, an idea that is supported by the observation that all cells in the primordium contribute fairly equally to directed migration^37^. We anticipate that this propulsion design likely drives the movement of other migrating tissues and should thus serve as a blueprint for cellular forces exerted by cells in other animals and contexts.

## Methods

### Zebrafish strains

Embryos were staged as previously described^38^. The *cxcl12a*^*at*30516^ allele contains a nonsense mutation resulting a premature stop codon^39^. Homozygous *cxcl12a*^*at*30516^ mutants were identified by the stalled migration of the primordium. The *lamC1*^*sa*9866^ allele was obtained from the Zebrafish International Resource Center (ZIRC, https://zebrafish.org) and contains a nonsense mutation resulting a premature stop codon^40^. Homozygous *lamC1*^*sa*9866^ mutants were identified by their shortened body axis^7^ or by PCR-based genotyping. Primers used for genotyping of the *lamC1*^*sa*9866^ mutation were the following; forward outer primer: 5’-TTCAGTTCATCGGGTTGC-3’, reverse outer primer: 5’-GTAACAGTTAGGGCACTGC-3’, forward inner primer: 5’-GTTTTCCTGCGTTGACGCTT-3’, reverse inner primer: 5’-GGTGTCGAGCGGTTGTAGAA-3’.

The PCR product was digested with the restriction enzyme BsaI (New England Biolabs, R0535L) to yield a 168 bp fragment for the wild-type allele and a 120 bp and a 48 bp fragments for the mutant allele. The *Tg(prim:mem-mCherry)*^41^, *Tg(cldnB:lyn*_2_*GFP)*^22^, *hsp70:cxcl12a*^42^, *TgBAC(cdh1:cdh1-TagRFP)*^29^, *TgBAC(cxcr4b:cxcr4b-Kate2-IRES-EGFP-CaaX)p7*^43^ and *TgBAC(cxcr4b:zGrad)*^29^ lines were previously described.

### Generation of mutant alleles

To generate *tln1*, *tln2a*, *tln2b*, *itgb1a* and *itgb1b* mutants, we followed previously described CRISPR-Cas9-based gene editing protocols^44^. mRNA for Cas9 was synthesized by *in vitro* transcription with mMESSAGE mMACHINE T7 Transcription kit (Thermo Fisher Scientific, cat no. AM1344) using the linearized plasmid *pST1374-NLS-flag-linker-Cas9* (Addgene 44758 ^45^) as a template. Three to four gRNAs were designed to target the coding sequence around the start codons in the cases where the genes were fully annotated in the Ensembl genome browser (GRCz10, www.ensembl.org). Otherwise, we designed gRNAs targeting the available coding sequence in the Ensembl genome browser (GRCz10). gRNA sequences were identified using the CRISPR guide design offered by Benchling (https://www.benchling.com/crispr/). The templates for gRNAs were synthesized by PCR. Briefly, a target sequence specific primer was designed which contained the T7 promoter sequence, the target sequence without the PAM site, and a overhang for primer annealing. A primer that coded for the chimeric gRNA backbone (sequences are shown below) was designed. All primers were purchased from Integrated DNA Technologies (IDT, https://www.idtdna.com). The target sequence specific primer and the chimeric gRNA backbone primer were annealed, filled-in by Taq polymerase and amplified by PCR. The PCR products were column-purified using the QIAquick PCR Purification Kit (Qiagen, 28106) and subjected to *in vitro* transcription using the MEGAscript T7 Transcription kit (Thermo Fisher Scientific, AM1334) to obtain the gRNAs. Three to four gRNAs (final concentration of each gRNA: 200 ng/ul) were mixed with Cas9 mRNA (300 ng/ul) and 1 nl was injected into one-cell stage embryos. The injected embryos were raised to adulthood and out-crossed to wild-type adults. Embryos from these crosses were genotyped for potential mutations induced by gene editing to identify adults with germ line mutations. Embryos from adults carrying germ line mutations were raised and genotyped as adults by PCR and sequencing. The target specific primers used and the mutant allele isolated for each gene are listed below.

The chimeric gRNA backbone primer: 5’-gttttagagctagaaatagcaagttaaaataaggctagtccgttatcaacttgaaaaagtggcaccgagtcggtgcggatc-3’

#### Generation and genotyping of the *tln1*^*d*4^ mutant

The following target sequence specific primers were used (*tln1* target sequences are underlined): gRNA1: 5’-TAATACGACTCACTATA<u>ACGACGCCTGTCGAATCATC</u>gttttagagctagaaatagcaag-3’ gRNA2:

5’-TAATACGACTCACTATA<u>ACAGGCGTCGTACACCACTG</u>gttttagagctagaaatagcaag-3’ gRNA3: 5’-TAATACGACTCACTATA<u>GGCGCTGTCGCTGAAGATCG</u>gttttagagctagaaatagcaag-3’ gRNA4: 5’-TAATACGACTCACTATA<u>TCTCTCCCTGATGATTCGAC</u>gttttagagctagaaatagcaag-3’

The isolated mutant fish harbor a 4 bp deletion in *tln1* exon 2 resulting in a frame shift that introduces a premature stop codon. *tln1*^*d*4^ mutant embryos recapitulate the previously described *tln1* phenotype such as partially penetrant cardiac edema and embryonic lethality^46^. Note that *tln1*^*d*4^ mutant fish can be kept as homozygous adults in the presence of the BAC transgene *TgBAC(tln1:tln1-YPet)* which rescues the lack of *tln1* activity. The *tln1*^A%^ allele was genotyped by amplifying the locus through PCR and digestion of the amplicon with the restriction enzyme Sau3AI (New England Biolabs, R0169L). The digest yields a 122 bp and a 53 bp fragment for the *tln1* wild-type allele and in a 171 bp fragment for the *tln1*^*d*4^ allele. The primers used for PCR were: forward outer primer: 5’-CAAGTGGCTCCGCCTGTACT-3’ reverse outer primer: 5’-ATAGGCCTAAAGGTATGCCAGC-3’ forward inner primer: 5’-GAGTAGCAGTGGCACAGTCC-3’ reverse inner primer: 5’-TGATGGACTCACGCTGGC-3’.

Note that the copy number of the *tln1* wild-type and the *tln1*^*d*4^ alleles with *TgBAC(tln1:tln1-YPet)* can be determined by the intensity of the bands of the digested amplicons on a 3% agarose gel with the above genotyping protocol.

#### Generation and genotyping of the *tln2a*^*i*23^ mutant

The following target sequence specific primers were used (*tln2a* target sequence is underlined): gRNA1: 5’-TAATACGACTCACTATA<u>CATGCCGGGTCATCAGAGAG</u>gttttagagctagaaatagcaag-3’ gRNA2: 5’-TAATACGACTCACTATA<u>TGTCTCTGAAGATCTGTGTG</u>gttttagagctagaaatagcaag-3’ gRNA3: 5’-TAATACGACTCACTATA<u>CATGCAGTTTGAGCCCTCAA</u>gttttagagctagaaatagcaag-3’ gRNA4: 5’-TAATACGACTCACTATA<u>GAACCCTCTCTCTGATGACC</u>gttttagagctagaaatagcaag-3’

The isolated *tln2a* mutant line comprises a 23 bp insertion (1 bp deletion plus 24 bp insertion) in *tln2a* exon 3 resulting in a frame shift that causes a premature stop codon. We were not able to obtain *tln2a*^*i*23^ homozygous adult fish in Mendelian ratios, suggesting that *tln2a*^*i*23^ homozygous mutant fish die at some point between 5 dpf and 60 dpf. However, we occasionally recovered *tln2a*^*i*23^ homozygous adult fish. The *tln2a*^*i*23^ allele was genotyped by PCR followed by SrfI restriction digest (New England Biolabs, R0629L). The primers used for PCR were: forward primer: 5’-CAGTTTGAGCCCTCAACGGCTGTATATGACGCATGCCcGGG-3’ reverse primer: 5’-CCCATATTCTGAAGCTGAGG-3’.

Note that the forward primer introduces a SrfI target site in the mutant allele only. Thus, SrfI restriction digest of the PCR amplicons results in a 192 bp fragment for the *tln2a* wild-type allele and a 37 bp and a 155 bp fragments for *tln2a*^*i*23^ mutant allele.

#### Generation and genotyping of the *tln2b*^*d*10^ mutant

The following target sequence specific primers were used (*tln2b* target sequence is underlined): gRNA1: 5’-TAATACGACTCACTATA<u>GGCCCCTGTGAGGGTCCTCG</u>gttttagagctagaaatagcaag-3’ gRNA2: 5’-TAATACGACTCACTATA<u>AGCCGCGAGGACCCTCACAG</u>gttttagagctagaaatagcaag-3’ gRNA3: 5’-TAATACGACTCACTATA<u>AGAACCTGATGGGAGCCGCG</u>gttttagagctagaaatagcaag-3’ gRNA4: 5’-TAATACGACTCACTATA<u>CAAGGGTGTGAAGCTGCTGG</u>gttttagagctagaaatagcaag-3’

The isolated mutant harbors a 10 bp deletion resulting in a frame shift that causes a premature stop codon. The predicted Tln2b^*d*10^ mutant protein comprises only the first 658 amino acids of the total 2580 amino acids. Note that *tln2b*^*d*10^ mutants can be kept as homozygous adult fish. The *tln2b*^*d*10^ allele was genotyped by PCR. The primers used for PCR were: forward outer primer: 5’-CAGGTGACCCCATAGACACG-3’ reverse outer primer: 5’-TGCATTGGTCACCTCTCCAG-3’ forward inner primer: 5’-TGTCCAAGGGTGTGAAGCTG-3’ reverse inner primer: 5’-CCTCTCCAGACGTGGGCTC-3’.

#### Generation and genotyping of the *itgb1a*^*d*34^ mutant

The following target sequence specific primers were used (*itgb1a* target sequence is underlined): gRNA1: 5’-TAATACGACTCACTATA<u>CCGGTGACCAACCGCAAGAA</u>gttttagagctagaaatagcaag-3’ gRNA2: 5’-TAATACGACTCACTATA<u>GAGAATCCTGAAGAATACAC</u>gttttagagctagaaatagcaag-3’ gRNA3: 5’-TAATACGACTCACTATA<u>GATAAGATCGAGAACCCGCA</u>gttttagagctagaaatagcaag-3’

The isolated mutant contains a 34 bp deletion in *itgb1a* exon 3 resulting in a frame shift that causes a premature stop codon. *itgb1a*^*d*34^ mutants can be kept as homozygous adult fish. The *itgb1a*^*d*34^ allele was genotyped by PCR. The following PCR primers were used: forward outer primer: 5’-GAGTTTCTGAAGCAGGGAG-3’ reverse outer primer: 5’-ATGGTGTTGCTTTCACACGC-3’ forward inner primer: 5’-AAAGAGGCTGCGCAGAAGAT-3’ reverse inner primer: 5’-TTTCTGAGGCTGGATCTGCG-3’.

#### Generation and genotyping of the *itgb1b*^*i*70^ mutant

The following target sequence specific primers were used (*itgb1b* target sequence is underlined): gRNA1: 5’-TAATACGACTCACTATA<u>CGTCATGCTCATGAGCTGAG</u>gttttagagctagaaatagcaag-3’ gRNA2: 5’-TAATACGACTCACTATA<u>TGTAAATGTTATCCTGCAGA</u>gttttagagctagaaatagcaag-3’ gRNA3: 5’-TAATACGACTCACTATA<u>GCATCGTGCTTCCTAATGAC</u>gttttagagctagaaatagcaag-3’

The isolated mutant contains a 70 bp insertion resulting in a frame shift that introduces two successive premature stop codons. The predicted mutant protein comprises only the first 320 amino acids of the total 806 amino acids. *itgb1b*^*i*70^ homozygous mutant embryos display a shorter body axis (Extended Data Fig. 5c–f) and die as embryos consistent with previous reports^47^. The *itgb1b*^*i*70^ allele was genotyped by PCR. The following PCR primers were used: forward outer primer: 5’-GATTGGACGCCGGGTATGTC-3’ reverse outer primer: 5’-AAACAGGCTGGAACTCCTCG-3’ forward inner primer: 5’-GGAATGTCACTCGTCTCC-3’ reverse inner primer: 5’-CATGGTGTAAATGTTATCCTGC-3’.

### Generation of transgenic lines

#### TgBAC(lamC1:lamC1-sfGFP)

For the *lamC1:lamC1-sfGFP transgene*, we used the BAC clone CHORI-211-194I4 which was obtained from BACPAC Resources, Children’ Hospital Oakland Research Institute, CA, USA. This BAC spans 192,491 bp of genomic DNA and contains the *lamC1* locus with about 50 kbp genomic sequence upstream of *lamC1* exon 1 and about 35 kbp genomic sequence downstream of *lamC1* exon 28. We modified this BAC clone in two ways by recombineering. First, the *tol2* sites and the *cryaa:Cerulean* transgenesis marker were inserted into the BAC backbone as described before^48^. Next, a cassette containing *sfGFP-FRT-galK-FRT*, flanked by 439 bp and 656 bp arms of homology upstream of the stop codon in *lamC1* exon 28 and downstream of the stop codon in *lamC1* exon 28, respectively, was inserted before the *lamC1* stop codon using galK-mediated recombineering^49^. The *galK* cassette was removed by Flippase-mediated recombination. The modified BAC clone was characterized in two ways. First, the BAC clone was digested with EcoRI (New England Biolabs, R0101L). Second, the modified locus was PCR-amplified and sequenced. This transgene expresses full length Laminin-*γ*1 fused to sfGFP from the *lamC1* promoter. The final BAC was purified with Nucleobond BAC 100 Kit (Takara Bio, 740579) and co-injected with 1 nl of 40 ng/ul *tol2* mRNA into one-cell stage embryos. The stable transgenic line was established by out-crossing the adult fish injected with the transgene and raising embryos with Laminin-*γ*1-sfGFP fluorescence and CFP fluorescence in the lenses of the eyes at 3 dpf. The full name of this transgenic line is *TgBAC(lamC1:lamC1-sfGFP)p1*. Note that this transgene recapitulates the previously reported expression pattern of *lamC1* mRNA expression^50^, partially rescues *lamC1* homozygous mutant embryos (Extended Data Fig. 2c) and does not affect primordium migration (Extended Data Fig. 2d).

#### TgBAC(tln1:tln1-YPet)

For the *tln1:tln1-YPet* transgene, we used the BAC clone DKEY-42J10 which was obtained from ImaGenes GmbH, Germany. The BAC clone spans 194,108 bp of genomic DNA. This includes 40 kb of genomic sequence upstream of the beginning of *tln1* exon 1, *tln1* exons 1 to 44, but lacks *tln1* exons 45 to 56. To include the complete coding sequence of *tln1* on the BAC we inserted the sequence of the missing *tln1* exons (exons 45 to 56) directly downstream of exon 44. This design was guided by the annotated *tln1* transcript *tln1-202* (ENSDART00000166799.2, Ensembl). We inserted the coding sequence for *YPet* between the head and the rod domains of *tln1*^51^, which is located in *tln1* exon 13. For this, we modified the BAC clone in four ways by recombineering. First, the *tol2* sites and the *cryaa:dsRed* transgenesis marker were inserted in the BAC backbone as previously described^48^. Second, a cassette comprising *tln1* exons 45-56 was PCR-amplified from cDNA of 1 dpf zebrafish embryos and the Kanamycin resistant gene *KanR*, flanked by 411 bp and 783 bp homology arms upstream of the end of exon 44 and downstream of the end of exon 44, respectively, was inserted directly after *tln1* exon 44. Third, a cassette consisting of *YPet-galK*, flanked by 175 bp and 327 bp homology arms to *tln1* exon 13, was inserted in exon 13. Fourth, the *galK* sequence was removed by seamless recombineering using an identical cassette to the cassette described above but lacking the *galK* sequence^49^. The amino acid sequence around the YPet insertion is Gly-Ser-Val-X-Ala-Leu-Pro, where YPet and restriction enzyme sequences were inserted in frame at position X. Each recombineering step was confirmed by EcoRI restriction digestion of the BAC clone and sequencing of PCR amplicons spanning the modified part on the BAC clone using primers outside the arms of homology. The final BAC was purified with Nucleobond BAC 100 Kit (Takara Bio, cat no. 740579) and co-injected with 1 nl of 40 ng/ul *tol2* mRNA into one-cell stage embryos. The stable transgenic line was established by out-crossing the adult fish injected with the transgene and raising embryos with Tln1-YPet expression and DsRed fluorescence in the lenses of the eyes scored at 3 dpf. The full name of this transgenic line is *TgBAC(tln1:tln1-YPet)p2*. Note that the expression pattern of this transgene recapitulates the *in situ* hybridization pattern against *tln1* mRNA (Extended Data Fig. 6a, d). This transgene rescued the lethality of *tln1* homozygous mutant embryos.

#### TgBAC(cxcr4b:F-tractin-mCherry)

For the *cxcr4b:F-tractin-mCherry* transgene, we used the BAC clone DKEY-169F10 which was obtained from ImaGenes GmbH, Germany. The BAC clone DKEY-169F10 contains a 60 kb genomic DNA fragment that spans the entire *cxcr4b* locus. First, the *tol2* sites and the *myl7:mScarlet* transgenesis marker were inserted in the BAC backbone as previously described^48^. The sequence of *F-tractin* (rat inositol 1,4,5-triphosphate 3-kinase A (ITPKA) amino acids 10-52 ^52^) together with a linker sequence (GLALPVAT) was assembled by primer annealing. A cassette consisting of *F-tractin-mCherry-FRT-galK-FRT*, flanked by homology arms 583 bp upstream from the beginning of *cxcr4b* exon 2 and 433 bp downstream of the *cxcr4b* stop codon, respectively, was inserted to replace the *cxcr4b* coding sequence in exon 2 (amino acids 6-358). The *galK* sequence was removed by Flippase-mediated recombination^49^. The modified BAC clone was characterized by EcoRI restriction digestion and sequencing of the PCR amplicon around the modified region. This transgene expresses the first five amino acids from *cxcr4b* exon 1 fused to F-tractin-mCherry from the *cxcr4b* promoter. The final BAC was purified with Nucleobond BAC 100 Kit (Takara Bio, 740579) and co-injected with 1 nl of 40 ng/ul *tol2* mRNA into one-cell stage embryos. The stable transgenic line was established by out-crossing the adult fish injected with the transgene and raising the embryos with F-tractin-mCherry expression and mScarlet fluorescence in the myocardium. The full name of this transgenic line is *TgBAC(cxcr4b:F-tractin-mCherry)p3*.

#### TgBAC(cxcr4b:F-tractin-mNeonGreen)

The *cxcr4b:F-tractin-mNeonGreen* transgenic line was generated as described above for the *cxcr4b:F-tractin-mCherry* line except that the BAC was modified to contain the *mNeonGreen* (mNG) instead of the *mCherry* coding sequence. The *mNG* coding sequence was PCR amplified from the Addgene plasmid 98886 ^53^.

### Generation of knock-in strains

#### itgb1b:itgb1b-sfGFP

The *itgb1b:itgb1b-sfGFP* knock-in line was generated as described previously^54^. First, we cloned part of the *itgb1b* genomic sequence spanning 29 bp of *itgb1b* exon 8 to the end of *itgb1b* exon 9 into the *pUC19* plasmid (exon numbering is based on transcript ID: ENSDART00000161711.2). Second, we inserted a linker sequence (amino acids GGPVAT), fused to the *sfGFP* coding sequence followed by the *SV40polyA* signal sequence before the stop codon of the *itgb1b* coding sequence in the *pUC19-itgb1b(exon8-exon9)* plasmid. A crRNA that targets *itgb1b* intron 8-9 (5’-GGAGGTCTTGATGTAGGATT-3’) was designed using the Custom Alt-R CRISPR-Cas9 guide RNA Design Tool (https://www.idtdna.com/pages/tools/alt-r-crispr-hdr-design-tool). The crRNA and the tracrRNA were purchased from IDT. Purified protein Cas9-NLS was purchased from QB3 MacroLab at UC Berkeley (https://macrolab.qb3.berkeley.edu/). The injection mix containing Cas9-NLS protein, crRNA, tracrRNA and the plasmid harboring the *itgb1b* targeting cassette was heat-activated and injected into one-cell stage wild-type embryos.

The injected embryos were raised to adulthood and out-crossed to wild-type adults. Embryos from these crosses were visually screened for Itgb1b-sfGFP fluorescence to identify the adults carrying the edited *itgb1b* locus. We isolated one *itgb1b:itgb1b-sfGFP* strain and confirmed the correct knock-in event by sequencing PCR amplicons spanning the genomic insertion site. Note that the expression pattern of *itgb1b:itgb1b-sfGFP* recapitulates the pattern seen by *in situ* hybridization against *itgb1b* mRNA (Extended Data Fig. 5a, i). *itgb1b:itgb1b-sfGFP* homozygous and *itgb1b:itgb1b-sfGFP/itgb1b*^*i*70^ trans-heterozygous adult fish are viable.

#### itgb1b:itgb1b^ΔNPxY^-sfGFP

The *itgb1b:itgb1b*^*ΔNPxY*^-*sfGFP* knock-in line was generated as described above for the *itgb1b:itgb1b-sfGFP* line except that the sequence coding for the first to the second NPxY motif (NPIYKSAVTTVVNPKY, amino acid 777-792) was deleted in the plasmid containing the targeting cassette. We identified one *itgb1b:itgb1b*^*ΔNPxY*^-*sfGFP* line and confirmed the correct knock-in event by sequencing PCR amplicons spanning the genomic insertion site.

### Generation of plasmid constructs

The *pDEST-tol2-acta1a-cxcl12a-t2a-mCherry* and *pDEST-tol2-acta1a-mCherry* plasmids were generated by Gibson cloning^55^ using the following plasmids as PCR templates: The plasmid backbone including the *tol2* sites, the *acta1a* promoter and the *cxcl12a* coding sequence were amplified from *pDestTol2pA2*^56^, *pDEST-tol2-acta1a-GFP*^57^ and *pCS2-cxcl12a*^58^, respectively. The *pCS2(+)-YPet-ZF1*, *pCS2(+)-mNeonGreen-ZF1* and *pCS2(+)-mCherry-ZF1* plasmids were generated by Gibson cloning using the following plasmids as templates: The plasmid backbone, the *YPet*, the *mNeonGreen* and *mCherry* sequences were amplified from *pCS2(+)-sfGFP-ZF1*^29^or *pCS2(+)*, *pUC19-tln1-5arm-YPet-3arm*, the Addgene plasmid 98886 ^53^ and *pDEST-tol2-hsp70l-secP-mCherry-SV40pA*^41^, respectively. The *pDEST- tol2-hsp70l-zGrad-t2a-mNeonGreen* plasmid was generated by Gibson cloning using the following plasmids as PCR templates: the plasmid backbone including *tol2* sites, the *hsp70l* promoter and the *zGrad* coding sequence were amplified from *pDEST-tol2-hsp70l-zGrad*^29^ and the *t2a-mNeonGreen* sequence was amplified from the Addgene plasmid 98886 ^53^ with PCR primers that also included the *t2a* sequence. The *pCS2(+)-lyn*_2_*mCherry* plasmid was generated based on the *pCS2(+)-lyn*_2_*EGFP* plasmid (gift from Reinhard W. Köster)^59^.

### *In vitro* mRNA transcription and mRNA injection

Linear templates for *in vitro* mRNA transcription were generated by restriction digest of plasmids or PCR on plasmids using primers containing a SP6 promoter sequence. mRNAs were transcribed using the mMESSAGE mMACHINE SP6 transcription Kit (Thermo Fisher Scientific, AM1340). Injection mixes contained 50 ng/*μ*l mRNAs with 0.1% Phenol Red Solution (Lifetechnologies, 15100-043). 1 nl of the injection mix was injected in one-cell-stage embryos. *tol2* mRNA was synthesized from a linearized *pCS2FA-transposase* plasmid^56^.

### Embryo dissociation and primary zebrafish cell culture

Glass bottom dishes (MatTek, P35G-0-20-C) were coated with mouse Laminin protein (Corning, 354232). Briefly, 50 *μ*g of mouse Laminin was mixed with Leibovitz L15 media without phenol red (Fisher, 21083027) and added to the glass bottom dishes. The dishes were incubated for 1 h at room temperature and then washed with PBS (137 mM NaCl, 2.7 mM KCl, 10 mM Na_2_HPO_4_, 1.8 mM KH_2_PO_4_). 33 hpf embryos were manually dechorionated and the tails dissected off in fish water (60 *μ*g/mL of Instant Ocean Sea Salts (Instant Ocean, SS15-10)) supplemented with 0.4 mg/ml MS-222 anesthetic (Sigma, A5040-25g). The tails were harvested and transferred into a cell dissociation medium (0.05% Trypsin-EDTA (Invtrogen, 25200-056)). The tails were then incubated in the dissociation medium at 28°C for 20–30 min, pipetted up and down with a P1000 pipette tip every 5 min until the tails were dissociated into single cells. This cell suspension was filtered through a 70 *μ*m nylon mesh (Fisher, 22-363-548). After a 3 to 4 washes with PBS, cells were resuspended in cell culture medium (Leibovitz L-15 medium without phenol red (Fisher, 21083027), 15% FBS (Fisher Scientific, cat no.A3160601), 100 U/ml Penicillin-Streptomycin (Invitrogen, cat no. 15140-122)). The cell suspension was added to the Laminin-coated dishes and incubated at 28°C overnight to allow the cells to settle.

### Immunofluorescence staining of cultured cells

Dissociated embryonic zebrafish cells seeded on Laminin-coated dishes were washed with PBS and fixed with 4% PFA in PBST at room temperature for 10 min. The cells were permeabilized with 0.5% Triton-X100 in PBS at room temperature for 10 min and blocked with 0.5% Bovine Serum Albumins (BSA, Millipore-Sigma, A4737-100G) in PBS at room temperature for 30 min, followed by incubation with the primary antibody overnight at room temperature. To detect Itgb1b-sfGFP and Tln1-YPet, we used the rabbit anti-GFP primary antibody (Torrey Pines Biolabs, NJ USA, cat no. TP401, lot no. 081211) at a dilution of 1:500 in 0.5% BSA/PBS. To detect F-tractin-mCherry, we used a sheep anti-mCherry primary antibody (custom made antibody by Covance^41^) at a dilution of 1:1000 in 0.5% BSA/PBS. The primary antibodies were detected with the following secondary antibodies at a 1:1000 dilution at room temperature for one hour: donkey anti-sheep-Cy3 (Jackson ImmunoResearch, PA USA, cat no. 713-166-147, lot no. 106361) and donkey anti-rabbit-Alexa488 (Jackson ImmunoResearch, PA USA, cat no. 711-546-152, lot no. 109010). The posterior lateral line cells were identified based on F-tractin-mCherry expression driven from the *cxcr4b* promoter. In the tails of 33 hpf embryos, the *cxcr4b* promoter drives high expression only in the posterior lateral line^29,58^. Imaging was performed using a spinning disk confocal Nikon W1 equipped with an Apo 60x NA 1.40 oil objective lens (Nikon, MRD01605). Images shown are maximum-projected z-stacks.

### Quantification of filamentous Collagen-IV by immunofluorescence staining

To stain for Collagen-IV and the membrane of the primordium cells in *lamC1* mutant and control embryos, we in-crossed *cldnB:lyn*_2_*GFP; lamC1+/−* fish and sorted for *cldnB:lyn*_2_*GFP; lamC1-/-* and *cldnB:lyn*_2_*GFP; lamC1−/+* or *cldnB:lyn*_2_*GFP; lamC1+/+* embryos at 30 hpf. One hour later, 31 hpf embryos were fixed with 4% PFA/PBST for 2 hrs at room temperature, then stored in 100% methanol (Millipore-Sigma, cat no. 322415-100ML) overnight at −20°C. Embryos were rehydrated using a series of 75%, 50% and 25% methanol in water and blocked in 1% BSA/PBST for 1 h at room temperature. The embryos were incubated in rabbit anti-collagen IV (1:200, ab6586, abcam, Cambridge UK) and goat anti-GFP (1:500, Covance, custom made antibody^43^) overnight at 4°C. Embryos were washed four times with PBST and incubated with donkey anti-rabbit Cy3 (1:500, Jackson ImmunoResearch, cat no. 711-165-152, lot no. 102215) and donkey anti-goat Alexa488 (1:500, Jackson ImmunoResearch, cat no. 705-546-147, lot no. 110667) secondary antibodies in PBST. Embryos were mounted in 0.5% low melt agarose/Ringer’s solution. Images were taken on a spinning disk confocal Nikon W1 microscope. The number of filament structures around the primordium was quantified using a custom-written macro in Fiji. Briefly, a 150 *μ*m × 150 *μ*m region of interest (ROI) containing the entire primordium was manually defined. The red fluorescent channel of the z-stack representing the Collagen-IV signal was sum-projected and only the fluorescence values above 1.25 the mean fluorescence intensity of the image were kept. Filamentous structures were extracted using the Tubeness filter (https://www.longair.net/edinburgh/imagej/tubeness/) in Fiji with sigma set to 1.0. Then, the image was thresholded using Otsu’s method and the number of filaments was counted with the Analyze Particles command in Fiji (settings: limiting size = 50-Infinity, circularity = 0.0-0.3).

### Whole mount *in situ* hybridization

The procedures for RNA probe synthesis and zebrafish embryo whole-mount *in situ* hybridization were performed as previously described^60^. The RNA probe against *cxcl12a* was previously described^58^. To synthesize the RNA probes against *tln1* and *tln2a*, parts of the transcripts were PCR-amplified from cDNA synthesized from zebrafish embryos using the primers indicated below and cloned into the *pCR2.1-TOPO* vector (Thermo Fisher Scientific, cat no. 451641). The plasmids were linearized using BamHI-HF (New England Biolabs, cat no. R3136L), column-purified (Qiagen, QIAquick PCR Purification Kit, cat no. 28106) and *in vitro* transcribed using a DIG RNA Labeling Kit (Roche, cat no. 11277073910) together with a SP6/T7 Transcription Kit (Roche, cat no. 10999644001). To synthesize the RNA probes against *tln2b*, *itgb1a*, *itgb1b*, *itgb1b.1*, *itgb1b.2*, *itgb2*, *itgb3a*, *itgb3b*, *itgb4*, *itgb5*, *itgb6* and *itgb7* DNA templates were PCR-amplified from cDNA synthesized from maternal, 28 hpf or 33 hpf embryonic cDNA using the primers listed below. The reverse primers harbor T7 promoter sequence at their a 5’ end so that the PCR products could directly be transcribed *in vitro* after column purification. The RNA probes were synthesized with the Roche DIG labeling mix (Roche, cat no. 11277073910) and detected using an anti-DIG antibody coupled to alkaline phosphatase (1:5000, Roche, cat no. 11093274910) and NBT/BCIP staining (Roche, cat no. 11681451001). Embryos were mounted on the 3% Methyl cellulose (Sigma Aldrich, cat no. M0512) and imaged on an Axioplan Microscope (Zeiss) equipped with an Axiocam (Zeiss) using a 10x (NA 0.5) objective lens.

Primer pairs:

*tln1*: forward: 5’-ATGGTACGGGGGCTGGAGAG-3’

reverse: 5’-ACCGCGCGAGCAGCAGCAGC-3’

*tln2a*: forward: 5’-TCCGGTATGTCAGGAGCAGC-3’

reverse: 5’-GGTTTCAACTGTCCCTCAGA-3’

*tln2b*: forward: 5’-TCGACTCCGCTCTCAGTGCT-3’

reverse: 5’-AATACTAATACGACTCACTATAGCACAAGCAGTTTCTTACTGG-3’

*itgb1a*: forward: 5’-GAAGCGGGAGAATCCAGAGG-3’

reverse: 5’-AATACTAATACGACTCACTATAGTCCATGGTCTTGACGACGTG-3’

*itgb1b*: forward: 5’-CCTACGTCTCCCACTGCAAG-3’

reverse: 5’-AATACTAATACGACTCACTATAGATTCGCACGTTCCACAAACG-3’

*itgb1b.1*: forward: 5’-AAAACCCCTGTTTTCCAAGCG-3’

reverse: 5’-AATACTAATACGACTCACTATAGCTCCGTTCTTGCAGTGGGAG-3’

*itgb1b.2*: forward: 5’-ATGTACTGAGCTTGACGGACG-3’

reverse: 5’-AATACTAATACGACTCACTATAGTGGACACACCATCAGGTAGC-3’

*itgb2*: forward: 5’-ATCCCCAAATCTGCAGTCGG-3’

reverse: 5’-AATACTAATACGACTCACTATAGCGTCGCATTCACAGTGTTCG-3’

*itgb3a*: forward: 5’-TCTGGGCAATAATCTGGCCG-3’

reverse: 5’-AATACTAATACGACTCACTATAGCGTGCCAACTGAAGGGTAGT-3’

*itgb3b*: forward: 5’-TCCAACCAGCAAAATGCACG-3’

reverse: 5’-AATACTAATACGACTCACTATAGATTTTGTCCTTGCACACGGC-3’

*itgb4*: forward: 5’-ACAATTTAGAATCGCGCTTCACC-3’

reverse: 5’-AATACTAATACGACTCACTATAGCGTTGGGTTTTCGGGGTTTC-3’

*itgb5*: forward: 5’-GTCACCCGCTGTGGAAGGATG-3’

reverse: 5’-AATACTAATACGACTCACTATAGATAGCGAGAGGTCCATGAGGTAG-3’

*itgb6*: forward: 5’-AAGATGCGCCTCCAGCTTAG-3’

reverse: 5’-AATACTAATACGACTCACTATAGGCTTCATGGAGTCGTTTCGC-3’

*itgb7*: forward: 5’-CATGTGCAGCTGTGACGAAG-3’

reverse: 5’-AATACTAATACGACTCACTATAGTCTCATGACAGGAGCCGCTAC-3’

*itgb8*: forward: 5’-GCCACCTAGAGGACAACGTC-3’

reverse: 5’-AATACTAATACGACTCACTATAGGTAGTGCAGGACGAGGGTTC-3’

### Dissection of embryos and collagenase treatment

The head and the yolk of 28 hpf wild-type embryos were removed in fish water supplemented with 0.4 mg/ml MS-222 anesthetic using forceps. Dissected embryos were transferred into deskin media (Ca^2+^ free Ringer’s solution, 50 mM EDTA, 0.4 mg/ml MS-222 anesthetic). The skin of the embryos was peeled off under a Leica dissection scope (Leica, Wild M420 with light stand) using forceps. The deskinned embryos were then transferred into Leibovitz L-15 media (Fisher, cat no. 11415064) supplemented with 0.4 mg/ml MS-222 anesthetic. The collagenase (Collagenase, Purified, 4 ku, Worthington, cat no. LS005275) stock was prepared in Leibovitz L-15 media at a concentration of 1000 U/ml. The deskinned embryos were soaked in the collagenase solution (909 U/ml) supplemented with 0.4 mg/ml MS-222 anesthetic at the room temperature for 30 min. After the treatment, the collagenase was washed out by Leibovitz L-15 media supplemented with 0.4 mg/ml MS-222 anesthetic, then the embryos were quickly mounted for atomic force or confocal microscopy.

### Atomic Force Microscopy measurements and data analysis

Deskinned embryos were glued to FluoroDish dishes (World Precision Instruments, FL USA, cat no. FD5040-100) using CELL-TAK (Corning, NY USA, cat no. 354240) and immersed in Leibovitz L-15 media supplemented with 0.4 mg/ml of the anesthetic MS-222 (Sigma Aldrich, cat no. A5040-25g). All Atomic Force Microscopy (AFM) measurements were carried out within 90 min after skin removal. The AFM measurements were performed on an Asylum Research MFP-3D-BIO Atomic Force Microscope using the Asylum Research software package Version IX (AR Software) as previously described^61^. The AR Software was used for cantilever calibration, force mapping, data export, and data visualization (Extended Data Fig. 3e). We used a spherical borosilicate glass bead probe with a 2.5 *μ*m radius, a spring constant 0.07 N/m, a Young’s modulus of 68.0 GPa, and a Poison ratio of 0.19 (Novascan, PT-GS). In contrast to pyramidal probes, which probe structures at the nm length scale such as extracellular filaments, spherical probes work at the *μ*m length scale and assess the global properties of the BM^62^. We used a trigger point of 1 nN force and an indentation velocity of 5.0 *μ*m/s. Such a slow indentation velocity minimizes viscous effects. Two to three two-dimensional 8-by-8 (20 *μ*m by 20 *μ*m) square grids were probed per deskinned embryo. The probed area was located above the muscles that overlie the notochord in the center of the deskinned embryo’s trunk (Extended Data Fig. 3d).

The AFM measurements were analyzed using the Rasylum package (https://github.com/nstone8/Rasylum), which runs on the R programming language software environment (https://www.r-project.org/). We modified the Rasylum package in three ways. First, we included a batch-mode option to analyze a set of force curves automatically. Second, we modified the extractStiffness function in the Rasylum package to specify the length of the force curve that will be fit to the Hertz model. The original extractStiffness function fits the entire force curve from the contact point to the maximum deformation point to the Hertz model. Our modification allows the user to only fit a select part of the curve to the Hertz model (from the contact point to a user specified length of the deformation part of the force curve, in our case 200 nm and 500 nm). Third, we modified the extractStiffness function to report statistical parameters for the fit of the probe approach part of the force curve (beginning of force curve to contact point) to a linear model and the indentation part of the force curve to the Hertz model (contact point to end of force curve/point of maximum deformation or the end point defined by the user). These statistical parameters were used as quality criteria to include or reject force curves. For the calculation of Young’s modulus, the sample Poisson ratio was assumed to be 0.45. The reduced Young’s modulus E* was obtained by fitting the first 200 nm of the approach curve past the contact point to the Hertz model

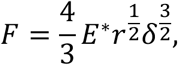

where *F* is the loading force, *E*\* is the reduced Young’s modulus, *r* is the radius of the spherical probe used, and *δ* is the indentation. The sample Young’s modulus *E*_*s*_ was calculated using

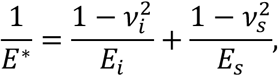

where *v*_*i*_ is the indenter’s Poisson ratio, *v*_*s*_ is the sample Poisson ratio, and *E*_*i*_ is the indenter’s Young’s modulus.

Due to debris after skin removal and the curvature of the BM above the muscle, some force curves were of low quality. To automatically select high quality force curves, we applied three criteria. The first criterion was the slope of the fit of the baseline of the approach curve (start of approach curve to contact point) to a linear equation. The second criterion was the residuals between the measured and fitted curve of the baseline of the approach curve to a linear equation. The third criterion was the P-value of the fit of the approach curve to the Hertz model from the contact point to 200 nm or 500 nm into the sample. Only force curves with a baseline slope between −10 to 10 pN/*μ*m, baseline sum-of-squared-residuals (SSR) smaller than 0.1 nN^2^, and P-values smaller than 1.0e-14 were included to determine the overall stiffness of the BM. These criteria select for force curves that have a flat baseline approach curve and a clearly defined contact point. Extended Data Fig. 3h shows representative force curves that meet and do not meet these criteria.

### Electron microscopy

*lamC1*-/- embryos were generated by in-crossing *lamC1*−/+ fish. 30 hpf embryos were fixed in EM fixative containing 2% paraformaldehyde, 2% glutaraldehyde in 0.1 M sodium cacodylate buffer at room temperature for 2 hrs and then overnight at 4°C. Fixed embryos were rinsed with 0.1 M sodium cacodylate buffer and post-fixed with 1% OsO_4_ in 0.1 M cacodylate buffer, followed by block-staining with 1% uranyl acetate aqueous solution overnight at 4°C. The samples were rinsed with water, dehydrated in graded series of ethanol, infiltrated with propylene oxide/Epon mixtures and finally embedded in EMbed812 (Electron Microscopy Sciences, PA USA). 70 nm sections were cut and mounted on 200 copper mesh grids and stained with uranyl acetate and lead citrate. Imaging was performed by Talos120C transmission electron microscope (Thermo Fisher Scientific, Hillsboro, OR) with Gatan (4k x 4k) OneView Camera (Gatan, Inc., Pleasanton, CA). The primordium cells were pseudo-colored using Adobe Illustrator 2020 (Adobe).

### Fluorescent recovery after photobleaching

All fluorescent recovery after photobleaching (FRAP) experiments were performed on a Nikon W1 spinning disk confocal microscope equipped with the Apo LWD 40X NA 1.15 objective lens (Nikon, cat no. MRD77410). 32 to 34 hpf embryos were mounted in 0.5% low melt agarose (National Diagnostics, cat no. EC-205)/Ringer’s solution on glass-bottom dishes, then immersed in Ringer’s solution supplemented with 0.4 mg/ml MS-222 anesthetic. The temperature of the mounted embryos was kept at 30°C using a Tokai Hit incubation system STXG-TIZWX-SET (Tokai Hit, Shizuoka Japan). For the analysis of Itgb1b-sfGFP at the myotendinous junctions, embryos were treated with 50 *μ*M Rockout (Rho Kinase Inhibitor III, MilliporeSigma, cat no. 555553-10MG) in 1% DMSO or control-treated with 1% DMSO for 3 hrs. The circular ROI with a 3 *μ*m radius was selected and bleached with a 405 nm laser. Images with a fixed z-position were taken every 0.5 sec for 30 sec or 1 min. Two regions of interest per embryo were recorded. For simultaneous FRAP of LamC1-sfGFP and sec-mCherry, 24 hpf *lamC1:lamC1-sfGFP*; *hsp70l:sec-mCherry* embryos were heat shocked at 39.5°C for 1 h to induce sec-mCherry expression from the heat shock promoter. To bleach LamC1-sfGFP and sec-mCherry, a circular ROI with a 6 *μ*m radius was selected and bleached with a 405 nm laser. Images with a fixed z-position were taken every 100 ms for 30 sec. For long-term LamC1-sfGFP FRAP, three to six circular ROI with a 6 *μ*m radius per embryo were selected and bleached with a 405 nm laser. Z-stack images were taken every 10 min for 1 h. All FRAP analyses were performed using the mean intensity values of the regions of interest extracted by the time-measurement setting in NIS-Elements (Nikon) except for the long-term LamC1-sfGFP FRAP. For the long-term LamC1-sfGFP FRAP, we first sum-projected the z-stacks and corrected for photo-bleaching using the mean fluorescent intensity of the image for each time point. The circular regions of interest with a 6 *μ*m radius were manually selected and tracked over time. The mean intensity in the regions of interest was then measured and plotted. For all other FRAP analyses, we first normalized the fluorescent intensity of the bleached region at each time point *I*_*x*_ to the difference between the fluorescent intensity before bleaching (*I*_max_) and the minimal fluorescent intensity after bleaching (*I*_min_) using

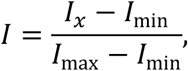

where *I* is the normalized fluorescence intensity. The fluorescent intensity in the bleached region was obtained from the FRAP module in NIS-Elements (Nikon). We then corrected for overall photo-bleaching by dividing the normalized fluorescent intensity *I* by the overall rate of photo-bleaching. To calculate the overall rate of photo-bleaching we randomly picked a 30 *μ*m × 30 *μ*m region outside of the bleached region and extracted the mean intensity over time from four movies for each experimental setting and averaged them. The overall photo-bleaching rate was calculated by dividing the average intensity of each time point by the averaged intensity of the first time point. The FRAP recovery curves were then fitted to an exponential plateau function in Prism (GraphPad)

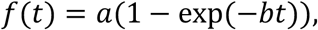

where *a* is the mobile fraction in percent and *b* is the rate constant of fluorescent recovery. The *t*_1/2_ was calculated using

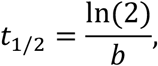

with *b* the rate constant of fluorescence recovery^63^. Our fluorescence recovery curves were measured for different lengths and we only fitted the first 28 sec to ensure comparability of the fits of different experiments.

### Primordium migration distance quantification

To quantify the cumulative migration distance of the primordium, the time-lapse image sequences were sum-projected and the tip of the primordium was tracked using the Manual Tracking plugin provided by Fabrice Cordelieres in Fiji (https://imagej.nih.gov/ij/plugins/manual-tracking.html).

### Ectopic expression of Cxcl12a from the trunk muscle cells

1 nl of 10 ng/*μ*l *pDEST-tol2-acta1a-cxcl12a-t2a-mCherry* plasmid DNA or *pDEST-tol2-acta1a-mCherry* plasmid DNA was injected together with 40 ng/*μ*l *tol2* mRNA into one-cell stage embryos obtained from the following crosses: *lamC1+/−; cxcl12a-/-; cldnB:lyn*_2_*GFP* in-cross, and *lamC1+/−; cxcl12a-/-; cxcr4b:cxcr4b-Kate2-IRES-GFP-CaaX* in-cross. *lamC1-/-* mutant embryos were identified by morphology. To score for the rescue of primordium migration through ectopic expression of Cxcl12a from muscle cells, the injected embryos were sorted at 32 hpf for *cldnB:lyn*_2_*GFP* and mounted in 0.5% low-melt agarose/Ringer’s solution supplemented with 0.4 mg/ml MS-222 anesthetic. Images were taken with a Leica 165M FC Fluorescent Stereo Microscope equipped with a Leica DFC345 FX camera (Leica Microsystems). The distance from the ear to the primordium was measured using Fiji. To observe receptor internalization induced by ectopic Cxcl12a secreted from the muscle cells, the embryos were mounted in the 0.5% low-melt agarose/Ringer’s solution supplemented with 0.4 mg/ml MS-222 anesthetic. 25–26 hpf embryos were imaged on Leica SP8 confocal microscope equipped with HyD detectors (Leica Microsystems) using a 40x (NA 1.1) objective. The power of the 488 nm and 561 nm laser lines was calibrated to 152 *μ*W and 110 *μ*W, respectively, using a power meter (X-Cite Power Meter Model, Lumen Dynamics, cat no. XR2100). The z-step size was set to 2.0 *μ*m. We manually cropped the z-stack such that it only contained the primordium. We then generated a binary mask based on the GFP channel and applied it to the RFP and GFP channels. The GFP and RFP channels were then sum projected, the mean intensity of each channel was measured and the RFP mean intensity was divided by the GFP mean intensity using a custom-written macro in Fiji.

### Local photo-bleaching of LamC1-sfGFP

All LamC1-sfGFP photo-bleaching experiments were performed on a spinning disk confocal microscope (Nikon W1) equipped with the Apo LWD 40X NA 1.15 objective lens (Nikon) and a 405 nm and a 473 nm laser line controlled by a miniscanner. The ROIs for bleaching were defined by a custom-written macro in Fiji and saved as a binary tiff image with 1024 x 1024 pixel resolution. The tiff image was imported into the Nikon NIS-Elements AR software as an ROI for the photo-bleaching experiments. The Fiji macro creates a hexagonal pattern with 20 x 20 dots (= 120 *μ*m × 120 *μ*m), each of which is separated from its neighboring dots by 6 *μ*m. A single dot was 2-pixel wide (= 0.65 *μ*m). Bleaching was performed with a 473 nm laser in a single plane.

### Live imaging

Extended live imaging (10 h) of the primordium in *itgb1b-/-; prim:mem-mCherry* embryos and control embryos was performed using a Leica SP8 confocal microscope. Embryos were generated by crossing *itgb1b:itgb1b-sfGFP/-; prim:mem-mCherry* fish to *itgb1b:itgb1b-sfGFP/-* fish. Mutants were identified by the absence of Itgb1b-sfGFP expression. Note that *itgb1b-/-* mutants produced by this cross showed a slightly stronger overall morphological defect than mutants generated by in-crossing *itgb1b−/+* fish, indicating that maternal Itgb1b rescues the lack of zygotic Itgb1b slightly better than maternal Itgb1b-sfGFP. For imaging embryos were mounted in the 0.5% low-melt agarose/Ringer’s solution supplemented with 0.4 mg/ml MS-222 anesthetic in the same dish. The time-lapse movies were taken on a Leica SP8 confocal microscope equipped with HyD detectors (Leica Microsystems) using a 20x (NA 0.5) objective and a heated stage (Warner Instruments, Quick Exchange Heated Base, cat no. QE-1). The following settings were used: z-step size 5.0 *μ*m, time interval 30 min, duration 9.5 h, temperature 28°C.

Short-term live imaging (1 h) of the migration of the primordium in *itgb1b-/-; cldnB:lyn*_2_*GFP* embryos and control embryos was performed using a spinning disk confocal microscope (Nikon W1) equipped with the Apo LWD 40X NA 1.15 objective lens (Nikon). 32 hpf embryos were mounted in 0.5% low melt agarose/Ringer’s solution on glass-bottom dishes, then immersed with Ringer’s solution supplemented with 0.4 mg/ml MS-222 anesthetic. The temperature of the medium was kept at 30°C using a Tokai Hit incubation system STXG-TIZWX-SET (Tokai Hit). *itgb1b-/-* embryos and control (sibling) embryos were sorted according to their body length before mounting. The body length of *itgb1b-/-* embryos is slightly shorter than *itgb1b−/+* or *itgb1b+/+* embryos. Images with a 40 *μ*m z-stack with a step size 0.8 *μ*m were taken every 5 min for 1 hour with a multi-position setting. Imaged embryos were digested and genotyped for *itgb1b* as described above. To image the deformation of the BM by skin basal cells, 32 hpf *TgBAC(lamC1:lamC1-sfGFP)* embryos injected with *lyn*_2_*-mCherry* mRNA at the 1-cell stage were mounted at 32 hpf in 0.5% low melt agarose/Ringer’s solution on glass-bottom dishes, and immersed with Ringer’s solution supplemented with 0.4 mg/ml MS-222 anesthetic. Bleaching and imaging was performed using a spinning disk confocal microscope (Nikon W1) equipped with the Apo LWD 40X NA 1.15 objective lens (Nikon). The temperature of the medium was kept at 30°C using a Tokai Hit incubation system STXG-TIZWX-SET (Tokai Hit). First, the LamC1-sfGFP-labeled BM was photo-bleached to generate a hexagonal dot pattern. Second, the location of the bleach pattern was imaged with the following parameters: z-step size 0.4 *μ*m, time interval 1 min, duration 10 min.

To image the deformation of the BM by the primordium, 32 hpf *TgBAC(lamC1:lamC1-sfGFP); prim:mem-mCherry* embryos were mounted in 0.5% low melt agarose/Ringer’s solution on glass-bottom dishes and immersed with Ringer’s solution supplemented with 0.4 mg/ml MS-222 anesthetic. Bleaching and imaging was performed using a spinning disk confocal microscope (Nikon W1) equipped with the Apo LWD 40X NA1.15 objective lens (Nikon). The temperature of the medium was kept at 30°C using a Tokai Hit incubation system STXG-TIZWX-SET (Tokai Hit). First, the LamC1-sfGFP-labeled BM was photo-bleached to generate a hexagonal dot pattern. Second, the location of the bleach pattern was imaged with the following parameters: z-step size 0.4 *μ*m, time interval 5 min, duration 120 min. The multi-position setting was used to image multiple embryos at the same time and in the same dish. The *TgBAC(lamC1:lamC1-sfGFP); prim:mem-mCherry; hsp70l:cxcl12a* control embryos were heat shocked at 39.5°C for 1 hour at 28 hpf to induce Cxcl12a expression and block primordium migration before imaging.

To image the accumulation of LamC1-sfGFP under the primordium, 32 hpf *TgBAC(lamC1:lamC1-sfGFP); TgBAC(cxcr4b:F-tractin-mCherry)* embryos were mounted in 0.5% low melt agarose/Ringer’s solution on glass-bottom dishes, then immersed with Ringer’s solution supplemented with 0.4 mg/ml MS-222 anesthetic. Imaging was performed using a spinning disk confocal microscope (Nikon W1) equipped with the Apo LWD 40X NA 1.15 objective lens (Nikon). The temperature of the medium was kept at 30°C using a Tokai Hit incubation system STXG-TIZWX-SET (Tokai Hit). The green and the red channels were imaged sequentially to prevent fluorescence bleed-through.

To image the localization of Itgb1b^*ΔNPxY*^-sfGFP in the primordium cells, 34 hpf *itgb1b:itgb1b*^*ΔNPxY*^*-sfGFP*; *prim:mem-mCherry* and *itgb1b:itgb1b-sfGFP*; *prim:mem-mCherry* control embryos were mounted in 0.5% low melt agarose/Ringer’s solution on glass-bottom dishes, then immersed with Ringer’s solution supplemented with 0.4 mg/ml MS-222 anesthetic. Images were taken on a Leica SP8 confocal microscope equipped with HyD detectors (Leica Microsystems) using a 40x (NA 1.1) objective and a z-step size of 0.42 *μ*m. The power of the 488 nm and 594 nm laser lines was calibrated to 95 *μ*W and 29 *μ*W, respectively, using a power meter (X-Cite Power Meter Model, Lumen Dynamics, cat no. XR2100). The GFP-to-mCherry fluorescence intensity ratio in the basal to apical axis was obtained by a custom-written macro in Fiji. Briefly, the macro generated a cell-membrane mask based on the mCherry channel using the Default thresholding method in Fiji. This mask was applied to the GFP and the mCherry channels and the signal intensities were obtained for each z-slices.

To assess the migration distance of the primordium in embryos in different genetic scenarios (*cldnB:lyn*_2_*GFP; lamC1-/-* and *cldnB:lyn*_2_*GFP* embryos with clones in muscle expressing mCherry from the *acta1a* promoter, *TgBAC(lamC1:lamC1-sfGFP)*), 26 hpf and 48 hpf embryos, respectively, were imaged on a Leica 165M FC Fluorescent Stereo Microscope equipped with a Leica DFC345 FX camera. The *TgBAC(lamC1:lamC1-sfGFP)* copy number was determined based on the intensity of the sfGFP fluorescence. Distance measurements were performed using Fiji.

The phenotype of *lamC1-/-; TgBAC(lamC1:lamC1-sfGFP)* embryos was documented using an Axioplan Microscope (Zeiss) equipped with an Axiocam (Zeiss) and a 5x (NA 0.25) objective. The *TgBAC(lamC1:lamC1-sfGFP)* was identified based on sfGFP fluorescence. The *lamC1* mutant embryos were identified and scored based on the morphological defects at 48 hpf.

The phenotype of *itgb1b-/-* embryos was documented on an Axioplan Microscope (Zeiss) equipped with an Axiocam (Zeiss) and a 5x (NA 0.25) objective. *itgb1b-/-* embryos were generated by crossing *itgb1b:itgb1b-sfGFP/-; prim:mem-mCherry* fish to *itgb1b:itgb1b-sfGFP/-* fish and *itgb1b-/-* embryos were identified by the absence of Itgb1b-sfGFP fluorescence.

To measure the migration distance of the primordium in *itgb1b-/-* embryos, 54 hpf embryos were generated by crossing *itgb1b:itgb1b-sfGFP/-; prim:mem-mCherry* with *itgb1b:itgb1b-sfGFP/-*. *itgb1b-/-* embryos were identified by the absence of Itgb1b-sfGFP expression. Images of embryos at 54 hpf were taken on a Leica 165M FC Fluorescent Stereo Microscope equipped with a Leica DFC345 FX camera. The distance was measured using Fiji. Note that the primordium reaches the tip of the tail by 42 hpf in wild-type embryos^38^.

To analyze the distance that the primordium migrated in various mutant backgrounds at 48 hpf, images were taken on a Leica 165M FC Fluorescent Stereo Microscope equipped with a Leica DFC345 FX camera. After imaging, the embryos were digested and genotyped if required. The following crosses were set up to obtain embryos of the indicated genotypes.

*(Z) tln1-/-* embryos: In-cross of *tln1-/-; TgBAC(tln1:tln1-YPet)/+; prim:mem-mCherry* fish, embryos were sorted against Tln1-YPet and for mem-mCherry.

*(Z) tln2a-/-* embryos: In-cross of *tln2a+/−; cldnB:lyn*_2_*GFP* fish, after the experiment embryos were genotyped for *tln2a*.

*(MZ) tln2b-/-* embryos: In-cross of *tln2b-/-; cldnB:lyn*_2_*GFP* fish.

*(Z) tln1-/-; (Z) tln2a-/-* embryos: In-cross of *tln1-/-; TgBAC(tln1:tln1-YPet)/+; tln2a+/−; prim:mem-mCherry*, embryos were sorted against Tln1-YPet and for mem-mCherry. After the experiment embryos were genotyped for *tln2a*.

*(Z) tln1-/-; (MZ) tln2b-/-* embryos: In-cross of *tln1-/-; TgBAC(tln1:tln1-YPet)/+; tln2b-/-; prim:mem-mCherry* fish, embryos were sorted against Tln1-YPet and for mem-mCherry.

*(Z) tln2a-/-; (MZ) tln2b-/-* embryos: In-cross of *tln2a+/−; tln2b-/-; cldnB:lyn*_2_*GFP* fish, after the experiment embryos were genotyped for *tln2a*.

*(Z) tln1-/-; (Z) tln2a-/-; (MZ) tln2b-/-* embryos: In-cross of *tln1-/-; TgBAC(tln1:tln1-YPet)/+; tln2a+/−; tln2b-/-; prim:mem-mCherry* fish, embryos were sorted against Tln1-YPet and for mem-mCherry. After the experiment, embryos were genotyped for *tln2a*.

*(MZ) itgb1a-/-* embryos: In-cross of *itgb1a-/-; cldnB:lyn*_2_*GFP* fish.

*(Z) itgb1b-/-* embryos: In-cross of *itgb1b−/+; cldnB:lyn*_2_*GFP* fish. After the experiment the embryos were genotyped for *itgb1b*.

*(MZ) itgb1a-/-; (Z) itgb1b−/+* embryos: Cross of *itgb1a-/-* female fish to *itgb1a−/+; itgb1b+/−; cldnB:lyn*_2_*GFP* male fish. After the experiment the embryos were genotyped for *itgb1a* and *itgb1b*.

*(Z) itgb1a−/+; (Z) itgb1b-/-* and *(Z) itgb1a-/-; (Z) itgb1b-/-* embryos: Cross of *itgb1a−/+; itgb1b:itgb1b-sfGFP/-; prim:mem-mCherry* fish to *itgb1a−/+; itgb1b:itgb1b-sfGFP/-* fish, embryos were sorted against Itgb1b-sfGFP and for mem-mCherry. After the experiment the embryos were genotyped for *itgb1a*.

### Circularity analysis of the primordium

To quantify the circularity of the primordium, we limited the length of the primordium to the first 100 *μ*m from the tip of the primordium. z-stacks were sum-projected. Using Fiji, the primordium region was manually cropped based on the intensity of Lyn_2_-GFP fluorescence. Then, a median filter with radius six pixel was applied and the background was subtracted using a rolling ball radius of 100 pixels. Images were rendered binary using the Huang thresholding algorithm to obtain a clear outline of the primordium. Finally, we quantified the circularity of the primordium for each time point using the ‘Analyze Particles’ macro in Fiji. The circularity *C* is defined such that a perfect circle yields a ratio of 1.

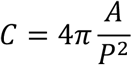

where *A* is the primordium’s area and *P* is the perimeter.

### Quantification of the LamC1-sfGFP intensity around the primordium

To quantify the lamC1-sfGFP intensity around the edge of the primordium and beyond the primordium, we used a semi-automated custom-written macro in Fiji. z-stacks were maximum-projected. The outer circumference of the primordium was manually traced using the primordium-specific mCherry signal as a guide. The encircled area was filled and thresholded. The thresholded image was duplicated. The first duplicate was 5 times eroded from its outline inward (=1.625 *μ*m). The second duplicate was dilated 5 times from its outline outward. Then the Two images were subtracted from each other. This generated a 10 pixel-wide (3.25 *μ*m) annulus-like area that we used as a binary mask. Next, the bright LamC1-sfGFP signal at the myotendinous junctions was thresholded using the LamC1-sfGFP signal and converted to a binary mask. This mask was dilated two times to remove the myotendinous junction signal from the analysis. These two masks were applied to the LamC1-sfGFP channel of the original image to extract the signal intensity of the LamC1-sfGFP around the primordium. To obtain the LamC1-sfGFP signal beyond the edge of the primordium (control measurement), the thresholded image of the primordium’s outer circumference was duplicated two times. The first and second duplicate was dilated 30 and 20 times, respectively, and subtracted from each other to generate a 10 pixel-wide (3.25 *μ*m) annulus-like area 6.5 *μ*m away from the primordium’s outer circumference as a binary mask. We plotted the mean intensity of the LamC1-sfGFP signal in each region from individual embryos.

### Analysis of actin flow

To image actin flow in single primordium cells, we co-injected BAC DNA coding for *cxcr4b:F-tractin-mNG* with 1 nl of 40 ng/ul *tol2* mRNA into one-cell stage wild-type and *itgb1b-/-* embryos also transgenic for *prim:mem-mCherry*. The *itgb1b-/-* embryos were generated by crossing *itgb1b:itgb1b-sfGFP/-; prim:mem-mCherry* to *itgb1b:itgb1b-sfGFP/-* and sorting for embryos lacking Itgb1b-sfGFP expression at 24 hpf. The embryos were mounted in the 0.5% low melt agarose/Ringer’s solution supplemented with 0.4 mg/ml MS-222 anesthetic at 32 to 34 hpf on a glass bottom dish and imaged on a spinning disk confocal microscope (Nikon W1) equipped with the SR HP Plan Apo 100X NA 1.45 objective lens (Nikon, MRD01905) at 30°C using a Tokai Hit incubation system STXG-TIZWX-SET. Clones in the primordium were identified based on mCherry and mNeonGreen expression and imaged on their basal side collecting single planes every 2 sec for 3 min. Actin flow analysis was performed by generating kymographs using Fiji. Briefly, a one pixel-wide 20 *μ*m ROI line was drawn manually from the cell’s center outward across the protrusion on the basal side. The kymograph was generated using the KymoResliceWide plugin provided by Eugene Katrukha and Laurie Young (https://imagej.net/KymoResliceWide). In singly labeled cells, the front of the actin flow and the front of the protrusion were identified visually, manually traced, and the actin flow rate and protrusion rate were extracted from the slopes of the trace lines. The actin polymerization rate was calculated by subtracting the actin flow rate from the protrusion rate.

### Whole-embryo imaging of transgenic expression patterns

The expression patterns of *TgBAC(lamC1:lamC1-sfGFP)*, *TgBAC(tln1:tln1-YPet)*, *itgb1b:itgb1b-sfGFP* and *itgb1b:itgb1b*^*ΔNPxY*^*-sfGFP* were imaged in 28 hpf embryos mounted in the 0.5% low melt agarose/Ringer’s solution supplemented with 0.4 mg/ml MS-222 anesthetic. The images were taken on a Leica SP8 confocal microscope equipped with HyD detectors (Leica Microsystems) using a 20x (NA 0.7) objective. The z-step size was set to 1.5 *μ*m. The images were stitched together using the auto tiling feature in the LAS X Life Science Microscope Software (Leica) and sum-projected.

### Blastomere transplantation and imaging of primordia with clones expressing Itgb1b-sfGFP and Tln1-YPet

We transplanted 20 to 50 cells from donor embryos at the 1000 to 8000-cell stage into host embryos of the same stage. All host embryos were wild type. Donor embryos were transgenic for *TgBAC(tln1:tln1-YPet)* or *itgb1b:itgb1b-sfGFP* and *Tg(prim:mem-mCherry)* or *TgBAC(cxcr4b:F-tractin-mCherry)*. At 28hpf, we isolated embryos that contained donor cells in the primordium based on the expression of the *Tg(prim:mem-mCherry)* or *TgBAC(cxcr4b:F-tractin-mCherry)* transgenes. Chimeric embryos were mounted in the 0.5% low melt agarose/Ringer’s solution supplemented with 0.4 mg/ml MS-222 anesthetic at 32-34 hpf on the glass bottom dish. Imaging was performed on a spinning disk confocal microscope (Nikon W1) equipped with the Apo LWD 40X NA 1.15 Water objective lens (Nikon) at 30°C using a Tokai Hit incubation system STXG-TIZWX-SET. Images were collected every 30 sec for 10 min as z-stacks with a z-step size of 1.0 *μ*m. The green and red fluorescent channels were sequentially scanned to prevent fluorescent bleed-through.

### Analysis of Itgb1b-sfGFP and Tln1-YPet localization in primordium cells

We analyzed the spatial distribution of Itgb1b-sfGFP and Tln1-YPet with respect to F-tractin-mCherry and membrane-tethered mCherry at the basal side of small clones of labeled primordium cells in two ways. First, we chose a single slice at the basal side of the cells in a clone. The clone contour was manually selected based on the F-tractin-mCherry or membrane-mCherry fluorescence using a 5 pixel-wide (= 1.625 *μ*m) segmented line to obtain the fluorescent intensity profile at each time point for the green and red channels in Fiji. Second, we assessed the co-localization of the fuorescently tagged proteins using the coloc2 plugin in Fiji. Before applying the coloc2 plugin, we processed the images as follows. Due to photo-bleaching only the first ten time points were included for analysis. Images were rotated such that the direction of migration aligned with the x-axis. Then the images were re-sliced to obtain transverse sections in the xz-plane. This resulted in images with the basal sides of the cells pointing upward (z-axis of imaging) and the direction of migration pointing to the right. Next, a median filter with 1 pixel width was applied and the images were rotated again such that the basal sides of the cells aligned horizontally. An ROI with a width of 3.25 *μ*m from the basal membrane inward was manually defined as the basal region of cells and used for the coloc2 analysis in Fiji with a custom-written macro. The macro compiles the individual ROIs of the basal region from each time point, creates a mask based on the sum intensities of the green and red channels using the Default method in Fiji. The individual ROIs were also compiled into a single image for each channel as a montage as shown in (Extended Data Fig. 7c). The macro then calculates the degree of co-localization, the Li’s ICQ value, using the coloc2 function in Fiji. Li’s ICQ is calculated as follows^64^. For each pixel in the ROI, the product of the difference of intensity and the mean intensity for each channel is calculated ((Ch1-mean(Ch1))*(Ch2-mean(Ch2)). Then, the number of pixels with a positive product are normalized to the total pixel number and 0.5 is subtracted. Therefore, Li’s ICQ ranges from −0.5 (signals perfectly segregated) to 0.5 (signals perfectly overlap). We performed this analysis for three consecutive xz-slices per clone, pooled them and showed in the panel.

### Blastomere transplantation and imaging of primordia with clones depleted in Talin activity

We transplanted 20 to 50 cells from donor embryos at the 1000 to 8000-cell stage into host embryos of the same stage. All host embryos carried *Tg(cldnB:lyn*_2_*GFP)*. Control donor embryos were transgenic for *Tg(prim:mem-mCherry)*. Talin-depleted donor embryos were obtained from an in-cross of *tln1-/-; tln2a+/−; tln2b-/-; TgBAC(tln1:tln1-YPet)/+; Tg(prim:mem-mCherry)* fish. Wild-type and Talin-depleted donor embryos were injected with 50 ng/*μ*l *zGrad* mRNA at the one-cell stage. 32-34 hpf embryos with chimeric primordia were mounted in 0.5% low melt agarose/Ringer’s solution supplemented with 0.4 mg/ml MS-222 anesthetic on a glass bottom dish. Imaging was performed on a spinning disk confocal microscope (Nikon W1) equipped with the Apo LWD 40X NA 1.15 Water objective lens (Nikon) at 30°C using a Tokai Hit incubation system STXG-TIZWX-SET. Images were collected every 5 min for 2 hours as Z-stacks with a Z-step size of 1.0 *μ*m. The green and red fluorescent channels were sequentially scanned to prevent fluorescent bleed-through. We analyzed three controls and three Talin-depleted chimeric primordia with donor cells at the primordium’s tip. Cumulative migration distance was quantified by manually tracking the tip of the primordia using the Manual Tracking plugin in Fiji. Kymographs were drawn using the KymoResliceWide plugin in Fiji.

### Generation of embryos lacking most Talin activity

Control wild-type embryos and embryos from an in-cross of *tln1-/-; tln2a+/−; tln2b-/-; TgBAC(tln1:tln1-YPet)/+; Tg(prim:mem-mCherry)* fish were injected with 50 ng/*μ*l *zGrad* mRNA at the one-cell stage. At 28 hpf, embryos were fixed with 4% PFA in PBST. After imaging, individual embryos were genotyped for *tln2a*.

### Temporal depletion of Talin activity by zGrad expression from the *hsp70l* promoter

Wild-type embryos and embryos from an in-cross of *tln1-/-; tln2a+/−; tln2b-/-; TgBAC(tln1:tln1-YPet)/+; Tg(prim:mem-mCherry)* fish were injected 1 nl of 5 ng/*μ*l *pDEST-tol2-hsp70l-zGrad-t2a-mNeonGreen* plasmid DNA together with 40 ng/*μ*l *tol2* mRNA at the one-cell stage. At 31 hpf, embryos were heat shocked at 39.5°C for 1 hour. For imaging embryos were mounted in the 0.5% low-melt agarose/Ringer’s solution supplemented with 0.4 mg/ml MS-222 anesthetic at 33 hpf. The time-lapse movies were taken from 34 hpf on a Leica SP8 confocal microscope equipped with HyD detectors (Leica Microsystems) using a 20x (NA 0.5) objective and a heated stage (Warner Instruments, Quick Exchange Heated Base, cat no. QE-1). The following settings were used: z-step size 3.0 *μ*m, time interval 20 min, duration 6 h, temperature 28°C. Only primordia with zGrad expressing clones marked by mNeonGreen expression were included in analysis. The cumulative migrated distance was analyzed as above.

### Analysis of skin-generated basement membrane wrinkles and traction

A 30 *μ*m × 30 *μ*m ROI centered on a transient increase in LamC1-sfGFP fluorescence intensity at the third time point in a 4-D stack spanning five time points with an 1 min interval was manually defined. The third point was set to 0 min. To visualize the LamC1-sfGFP increase as a graph, the stack was max-projected along the z-axis. The fluorescent intensity profile was extracted along a 10 *μ*m line ROI across the LamC1-sdfGFP increase at the 0 min time point. The same fluorescent intensity profiles along the same line ROI were obtained for the −1 min and +1 min time points. The fluorescence intensity was normalized to the mean of the −1 min time point. Traction was analyzed using Embryogram, and calculated and visualized with ParaView (ParaView-5.8.1, Kitware). For the traction stress calculation a Young’s modulus of 566.7 Pa was used for the BM based on the AFM measurements (Fig. 2d) and a Poisson ratio of 0.45. Traction was obtained using the −2 min time point as a reference for the undeformed BM. To quantify the temporal change of traction stresses around the local increases in LamC1-sfGFP, we averaged the traction stresses measured at the three marks closest to the LamC1-sfGFP increase. Particle Image Velocimetry (PIV) analysis of the basal skin cells was performed for detecting the displacement of the membrane from −1 min to 0 min using the PIVlab plugin (version 2.38 by William Thielicke) in MATLAB (Version 9.9, MathWorks). Every second vector was visualized on the cell membrane image at the −1 min time point. The displacement of the BM was obtained by Embryogram. The 3D quiver plots of the displacement of the BM from −2 min (reference) to 0 min with the scale factor 3 were drawn using ParaView (ParaView-5.8.1, Kitware).

### Analysis of primordium-generated traction stresses and angles of BM deformation

Four-dimensional confocal z-stacks were denoised by applying a median filer (width 2 pixels) and analyzed in Embryogram. In Embryogram, an area containing a well-defined bleach pattern that was in front of the primordium at the first time point of the time lapse was manually selected. Bleached cylinders with radii between 2–8 pixels (=0.65–2.6 *μ*m) were identified and tracked over time in the time lapse and manually curated in Embryogram. To subtract rigid motions, we manually selected cylinders to calculate the global displacement. If the bleach pattern spanned the horizontal myoseptum and extended past the primordium on either side, we used the two rows of bleached cylinders furthest from the horizontal myoseptum on both sides of the bleach pattern. If the bleach pattern did not span the horizontal myoseptum, we used the two rows of bleached cylinders furthest from the horizontal myoseptum on one side only. To perform the Finite Element Analysis, we constructed volumetric meshes on both sides of the BM to represent the skin and the muscle above and below the primordium, used up-sampling of 2, discretization order of 2, a Young’s modulus of 566.7 Pa and a Poisson ratio of 0.45. We excluded the cylinders on the edge of the bleach pattern from the analysis because these cylinders were often tracked incorrectly.

To analyze the angles between the displacement vectors and the direction of migration, and the traction stresses under the primordium, we exported data files containing 1) the X, Y and Z coordinates, 2) the displacement vector, and 3) the traction stress vector for each extrapolated vertex from ParaView (ParaView-5.8.1, Kitware). Then, we semi-manually selected vertices in front, middle, and rear part of the primordia defined as the 0–15 *μ*m, 15–65 *μ*m and 65 < from the tip of the primordium, respectively. For this analysis, we used the 40, 60 and 80 min time points of the time lapse videos. We also included cylinders with a distance of less than 10 *μ*m to the primordium in the analysis. For controls, we used cylinders that are at a distance less than 20 *μ*m from the horizontal myoseptum on the dorsal and ventral sides along the migratory route of the primordium. This corresponds to the maximal width of the primordium. In Fiji, we maximum-projected the primordium channel, manually traced the primordium edge, converted it to a binary mask and dilated the mask 10 *μ*m. Then, we applied this binary mask to the maximum-projected BM channel at the first time point. The approximate coordinates of the cylinders within the mask were manually recorded in Fiji. Using a custom-written R script, we compared these approximate cylinder-coordinates to the actual cylinder-coordinates obtained from the tracking in Embryogram by minimizing the distance between these two, and extracted information of actual cylinders only in the front, middle and rear of the primordium. BM displacement angles were analyzed based on the displacement of the bleached cylinder in the XY-plane in relation to the direction of primordium migration or the horizontal myoseptum in the case of the controls. The direction of primordium migration/horizontal myoseptum was obtained manually in Fiji. Half-circle polar diagrams were drawn using MATLAB. The cosines of these angles are shown in Fig. 2k. The magnitude of traction stress in the X- and Y-directions and in the Z-direction were extracted from cylinder located as described above.

3-dimensional and 2-dimensional displacement vectors, the magnitude of the traction stresses, the components of the stress tensors, and the magnitude of the stresses in the direction of primordium migration/horizontal myoseptum were visualized using ParaView (ParaView-5.8.1, Kitware). To calculate the stress in the direction of primordium migration/horizontal myoseptum, we manually obtained three dimensional vectors pointing in the direction of primordium migration/horizontal myoseptum in Fiji and extracted the stress using the calculator function in ParaView. The confocal stack of the primordium channel was binarized using the Li’s thresholding method in Fiji and superimposed. The data were visualized in ParaView.

## Acknowledgments

We thank R. Lehmann, L. Christiaen, D. Rifkin, M. Schober, J. Torres-Vázquez, W. Qian, P. Vagni, S. Lau and T. Colak-Champollion for critical comments, T. Gerson, T. Colak-Champollion and A. Feitzinger for reagents, T. Gerson, J. Proietti and S. Pirani for excellent fish care, N. Paknejad for advice on AFM, M. Cammer and Y. Deng for advice on microscopy, A. Liang, C. Petzold and K. Dancel-Manning for consultation and assistance with TEM work, and A. Ferrari and N. Chala for AFM consultation. The use of the NYULH DART Microscopy Laboratory (P30CA016087) and the Memorial Sloan Kettering Molecular Cytology Core Facility (P30 CA008748) is gratefully acknowledged. For providing the zebrafish knockout allele *lamC1*^*sa*9866^, we thank the Zebrafish International Resource Center (ZIRC). This work was supported by NIH grant NS102322 (H.K.), by an NYSTEM fellowship C322560GG (N.Y.), by an American Heart Association fellowship 20PRE35180164 (N.Y.), in part through the NYU IT High Performance Computing resources, services, and staff expertise, the NSF CAREER award 1652515 (D.N.), the NSF grants IIS-1320635 (D.N.), OAC-1835712 (D.N.), OIA-1937043 (D.N.), CHS-1908767 (D.N.), CHS-1901091 (D.N.), a gift from Adobe Research (D.N.), a gift from nTopology (D.N.), and a gift from Advanced Micro Devices, Inc (D.N.).

## Author contributions

N.Y., D.P., and H.K. conceptualized the study and designed the experiments. N.Y. performed all the zebrafish experiments with support from H.K. except for the AFM measurements which were performed by B.W. with samples prepared by N.Y. Embryogram software was developed by Z.Z., T.S. and D.P. with inputs from N.Y. and H.K. N.Y. analyzed most of the data with help from Z.Z. and T.S. for the traction stress analysis and from Z.Z. and B.W. for the AFM data analysis. N.Y. and H.K. wrote the main manuscript with input from Z.Z., T.S. and D.P. Z.Z., T.S. and D.P. wrote Supplemental Note with input from N.Y. and H.K. All authors approved of and contributed to the final version of the manuscript.

## Competing interests

No competing interests declared.

## Notes

### Competing Interest Statement

The authors have declared no competing interest.

